# Heterogeneous endocrine cell composition defines human islet functional phenotypes

**DOI:** 10.1101/2024.11.20.623809

**Authors:** Carmella Evans-Molina, Yasminye D. Pettway, Diane C. Saunders, Seth A. Sharp, Thomas SR. Bate, Han Sun, Heather Durai, Shaojun Mei, Anastasia Coldren, Corey Davis, Conrad V. Reihsmann, Alexander L. Hopkirk, Jay Taylor, Amber Bradley, Radhika Aramandla, Greg Poffenberger, Adel Eskaros, Regina Jenkins, Danni Shi, Ke Xu, Hakmook Kang, Varsha Rajesh, Swaraj Thaman, Fan Feng, Jean-Philippe Cartailler, Alvin C. Powers, Kristin Abraham, Anna L. Gloyn, Joyce C. Niland, Marcela Brissova, the Integrated Islet Distribution Program

## Abstract

Phenotyping and genotyping initiatives within the Integrated Islet Distribution Program (IIDP), the largest source of human islets for research in the U.S., provide standardized assessment of islet preparations distributed to researchers, enabling the integration of multiple data types. Data from islets of the first 299 organ donors without diabetes, analyzed using this pipeline, highlights substantial heterogeneity in islet cell composition associated with hormone secretory traits, sex, reported race and ethnicity, genetically predicted ancestry, and genetic risk for type 2 diabetes (T2D). While α and β cell composition influenced insulin and glucagon secretory traits, the abundance of δ cells showed the strongest association with insulin secretion and was also associated with the genetic risk score (GRS) for T2D. These findings have important implications for understanding mechanisms underlying diabetes heterogeneity and islet dysfunction and may provide insight into strategies for personalized medicine and β cell replacement therapy.

## INTRODUCTION

Diabetes mellitus impacts over 11% of Americans and is the eighth leading cause of death in the U.S., with an estimated economic cost of over $413 billion annually^1,2^. Type 1 diabetes (T1D) accounts for 5-10% of all diabetes cases and results from autoimmune-mediated destruction and loss of the insulin-producing β cells, whereas type 2 diabetes (T2D), the most prevalent diabetes form, is characterized by β cell dysfunction and peripheral insulin resistance. While the etiologies of T1D and T2D are largely distinct, both forms require loss and/or dysfunction of pancreatic β cells, have a strong genetic component, and are heterogeneous regarding disease pathophysiology, progression, and response to therapeutic interventions ^3–7^. Increasingly, data from clinical cohorts, tissues derived from human organ donors and mouse models highlight a role for impaired function of other islet cell subtypes, including α cells and δ abnormalities in diabetes pathophysiology^8–18^. Murine models have advanced our understanding of disease pathophysiology; however, there are critical differences in the function and architecture of murine and human pancreatic islets necessitating the comprehensive study of human islets to translate basic science research findings^19–25^.

The Integrated Islet Distribution Program (IIDP), formerly the Islet Cell Resource^26^, has served as the main source of human islets for research within the U.S., with nearly 300 million islets supplied for use in more than 600 unique studies to date (https://iidp.coh.org/). Located at City of Hope (Duarte, CA) and funded by the National Institutes of Diabetes and Digestive and Kidney Diseases (NIDDK) with targeted programmatic funding from Breakthrough T1D (formerly JDRF), the IIDP’s mission is to distribute high-quality human islets to the diabetes research community to support basic science and translational research. The IIDP subcontracts with a group of expert islet isolation centers at institutions across the country, who procure donated pancreata from their local Organ Procurement Organizations (OPOs). Rigorous protocols are followed to isolate the islets while maximizing the yield, viability and purity. An automated algorithm developed by the IIDP ensures fair and equitable islet distribution, matching the islet offers to awaiting researchers who have subscribed to receive islets and other non-islet biospecimens from the IIDP^27,28^.

In response to the NIH’s call to enhance rigor, reproducibility, and transparency in biomedical research, the IIDP initiated the Human Islet Phenotyping Program (HIPP) at Vanderbilt University Medical Center and the Human Islet Genotyping Initiative (HIGI) Program at Stanford University to provide systematic phenotypic and genotypic assessment of IIDP-supported human islet preparations (**Extended Data Figure 1**)^29^. This workflow comprehensively evaluates islet morphology, purity, and viability (**Extended Data Figure 2**). For the first time, it also assesses islet cell composition and concurrently examines the *in vitro* β and α cell function^12,30–35^. It also generates the genetic risk scores (GRS) for T1D and T2D while predicting genetic ancestry across a diverse group of organ donors in the U.S. (see **Figure 1A**). This centralized approach reduces redundancy and provides cost savings to investigators to accelerate the speed of scientific discoveries made using these islets. Importantly, the IIDP provides this rich integrated islet data generated by the HIPP and HIGI for researchers to explore via the online IIDP Research Data Repository (RDR). Here, we present an analysis of the first 299 IIDP-supported islet preparations from donors without a clinical diagnosis of diabetes analyzed using the HIPP and HIGI pipelines. Our integrated analysis of this dataset highlights marked heterogeneity in islet secretory traits and points to a previously underappreciated heterogeneity in islet cell composition. We found that the heterogeneity concerning three primary endocrine cell types, α, β, and δ cells, is associated not only with islet hormone secretion traits but also with sex, reported race and ethnicity, genetically predicted ancestry, and T2D GRS. While α and β cell composition influenced insulin and glucagon secretory traits, most notably, the abundance of a relatively small δ cell population showed the strongest association with insulin and glucagon secretion traits. Delta cell composition was also associated with the T2D GRS. These findings have important implications for understanding the mechanisms underlying personalized signatures—whether genetic, functional, or architectural— that predispose individuals to diabetes. Additionally, they may influence the progression and treatment outcomes of the disease, as well as inform β cell replacement therapy using human islets for transplantation.

**Figure 1.**
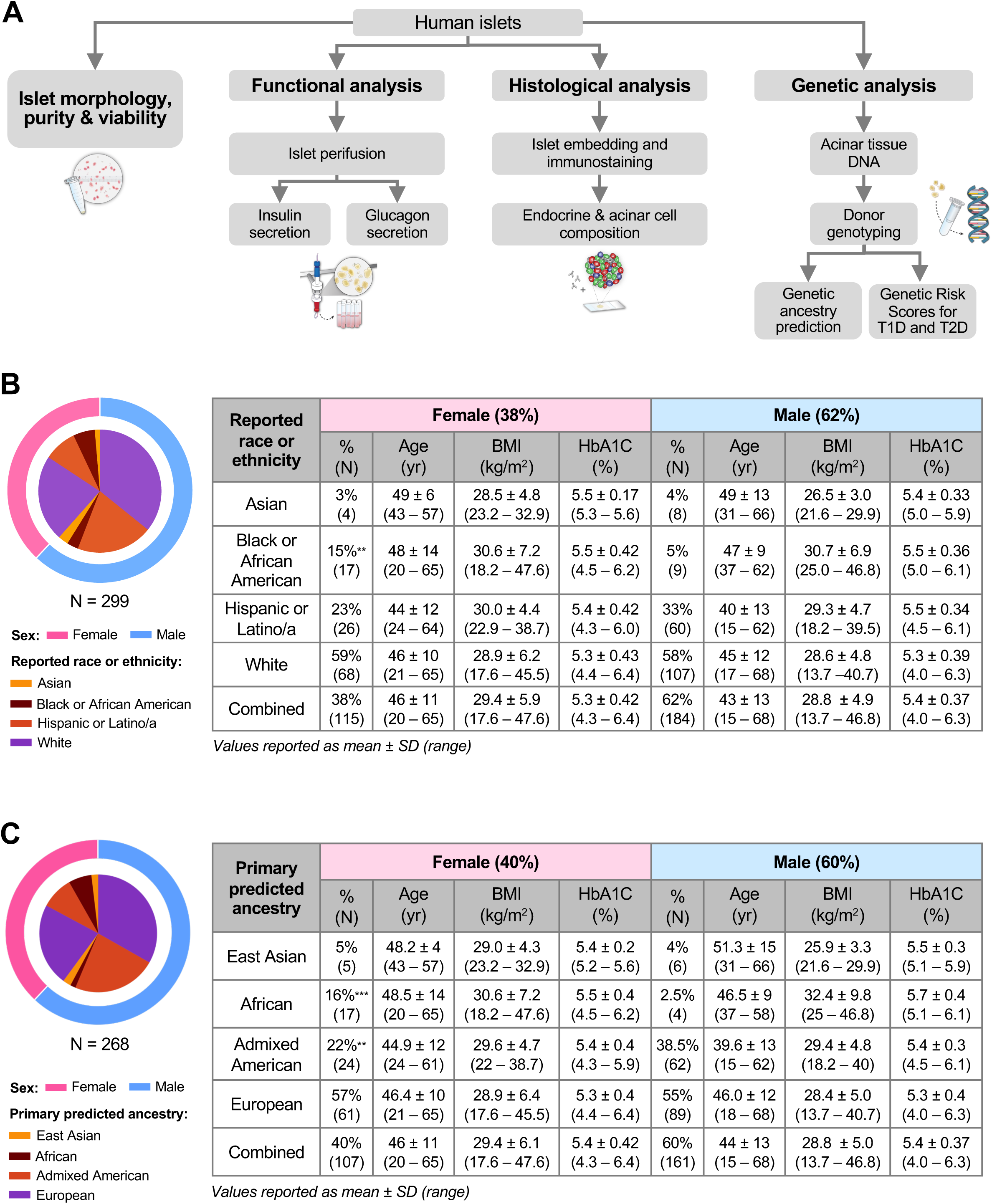
Integration of phenotyping and genotyping assessment of a diverse set of IIDP-derived human islet preparations. (**A**) Schematic illustrating the human islet phenotyping and genotyping pipelines utilized by the HIPP and HIGI. The HIPP performs analysis of islet morphology, purity, viability, histology, and function; the remaining sample is archived in a biorepository for additional analyses. The HIGI receives a non-islet tissue sample and isolates DNA to sequence for genetic ancestry prediction and calculation of genetic risk scores for type 1 and type 2 diabetes. (**B**) Descriptive statistics of donor demographics by sex and reported race or ethnicity (n = 299). Values are reported as mean ± standard deviation, followed by the range. (**C**) Descriptive statistics of donor demographics by sex and primary predicted ancestry (n = 268). Values are reported as mean ± standard deviation, followed by the range. **p < 0.01 vs. male donors; ***p < 0.001 vs. male donors.

## RESULTS

We present an analysis of the first release of 299 human islet preparations from donors without diabetes, for which all relevant donor information was available (**Figure 1B**). This dataset was derived from a diverse pool of donors with respect to reported race, ethnicity, age, and BMI. Islets from male donors were overrepresented in the dataset (62%), which is reflective of overall trends in available islets and organ donors in the U.S.^36^. The distribution of reported race or ethnicity differed between male and female donors (p = 0.014), where the dataset included a higher percentage of female donors that were reported Black or African American (15% vs. 5%, p = 0.006). A higher percentage of male donors were Hispanic or Latino, though this difference was not statistically significant). In 90% of the human islet preparations for which genotyping data was available, the genetic ancestry of the donor was predicted (n = 268; **Figure 1C**). Overall, there was a high concordance between reported race or ethnicity and primary predicted genetic ancestry within the two datasets (κ= 0.867, p < 0.001). Similar to the larger dataset, there was a lower percentage of male donors with primary African ancestry (2.5% vs. 16%, p = 0.0002) and a higher proportion of male donors with Admixed American ancestry (38.5% vs. 22%, p = 0.009).

Demographic features were similar in the group of donors where genetic ancestry was predicted (**Figure 1B-C**). There were no significant differences in age, BMI, or HbA1c between male and female donors by Wilson rank sum test. The mean age of male and female donors was 43 (range 15 – 68 years) and 46 years (range 20 – 65 years), and BMI averaged 29 kg/m^2^ in both male (range 13.7 – 46.8 kg/m^2^) in female donors (range 17.6 – 47.6 kg/m^2^). The mean HbA1c of male and female donors was 5.4% (range 4 – 6.3%) and 5.3% (range 4.3 – 6.4%). Of note, this dataset includes islets from donors within a normal HbA1c range, defined by the American Diabetes Association as <5.7% (n = 223, mean ± SD: 5.2 ± 0.3, range 4.0 – 5.6), as well as a subset of islets from donors with an HbA1c ≥ 5.7% (n = 76, mean ± SD: 5.9 ± 0.2, range 5.7 – 6.4). Together, these data form a powerful dataset to understand population-based differences in human islet phenotype.

### Islet processing traits associated with islet purity and viability

IIDP-affiliated islet isolation centers identify, process donor pancreata, and distribute human islet preparations for phenotypic assessment by the HIPP, genotypic assessment by the HIGI, and scientific investigation by researchers (**Supplemental File 1**). The IIDP RDR includes data on islet purity and viability, which are measured by islet isolation centers and made available to researchers at broadcast (i.e., prior to distribution, at the time the islets are offered to investigators). The post-shipment islet purity and viability is assessed by the HIPP on the day of arrival. The RDR also includes data related to islet processing, including organ cold ischemia duration, pre-shipment culture time, and transit time. The current dataset included human islet preparations from five islet isolation centers: Scharp-Lacy Research Institute (n = 113), Southern California Islet Cell Resource Center (n = 94), University of Miami Diabetes Research Institute (n = 30), University of Pennsylvania Islet Transplant Center (n = 21), and the University of Wisconsin Human Islet Core (n = 35). Pre-shipment culture time averaged 57.5 hours (range 8 – 237 hours), and the average transit time to the HIPP was 25 hours (range 10 – 102 hours). The mean islet purity measured by the HIPP was 73% (range 15 – 95%), and the mean dispersed islet cell viability was 76% (range 43 – 93%). Islet transit time and dispersed cell viability were positively correlated (Spearman r =0.2, unadjusted p-value (p_unadj_) = 0.001, **Extended Data Figure 3**). The HIPP assessment of islet purity did not show an association with cold ischemia duration, pre-shipment culture time, or islet transit time.

### Islet secretory function is highly heterogeneous amongst donors

A dynamic perifusion system was used to investigate simultaneously β and α cell hormone secretion in response to both physiologic and pharmacologic stimuli with co-stimulatory and opposing effects on insulin and glucagon secretion. This included basal glucose (5.6 mM), high glucose (16.7 mM), high glucose with isobutylmethylxanthine (IBMX), low glucose (1.7 mM) with adrenaline (Ad), and potassium chloride (KCl). On average, high glucose led to a 7.9-fold increase in stimulated insulin secretion and inhibited glucagon secretion by 0.4-fold on average compared to basal glucose. In contrast, low glucose with adrenaline led to the opposite effect, reducing insulin secretion by approximately 0.2-fold and increasing glucagon secretion by 5.1-fold compared to basal glucose. Exposure to IBMX, a phosphodiesterase inhibitor that increases intracellular cAMP levels, potentiated glucose-stimulated insulin secretion (14-fold) and stimulated glucagon secretion (6.5-fold) compared to basal glucose level. Additionally, KCl-induced membrane depolarization potently stimulated both insulin and glucagon secretion by approximately 31– and 11-fold on average, respectively (**Figure 2A-B**).

**Figure 2.**
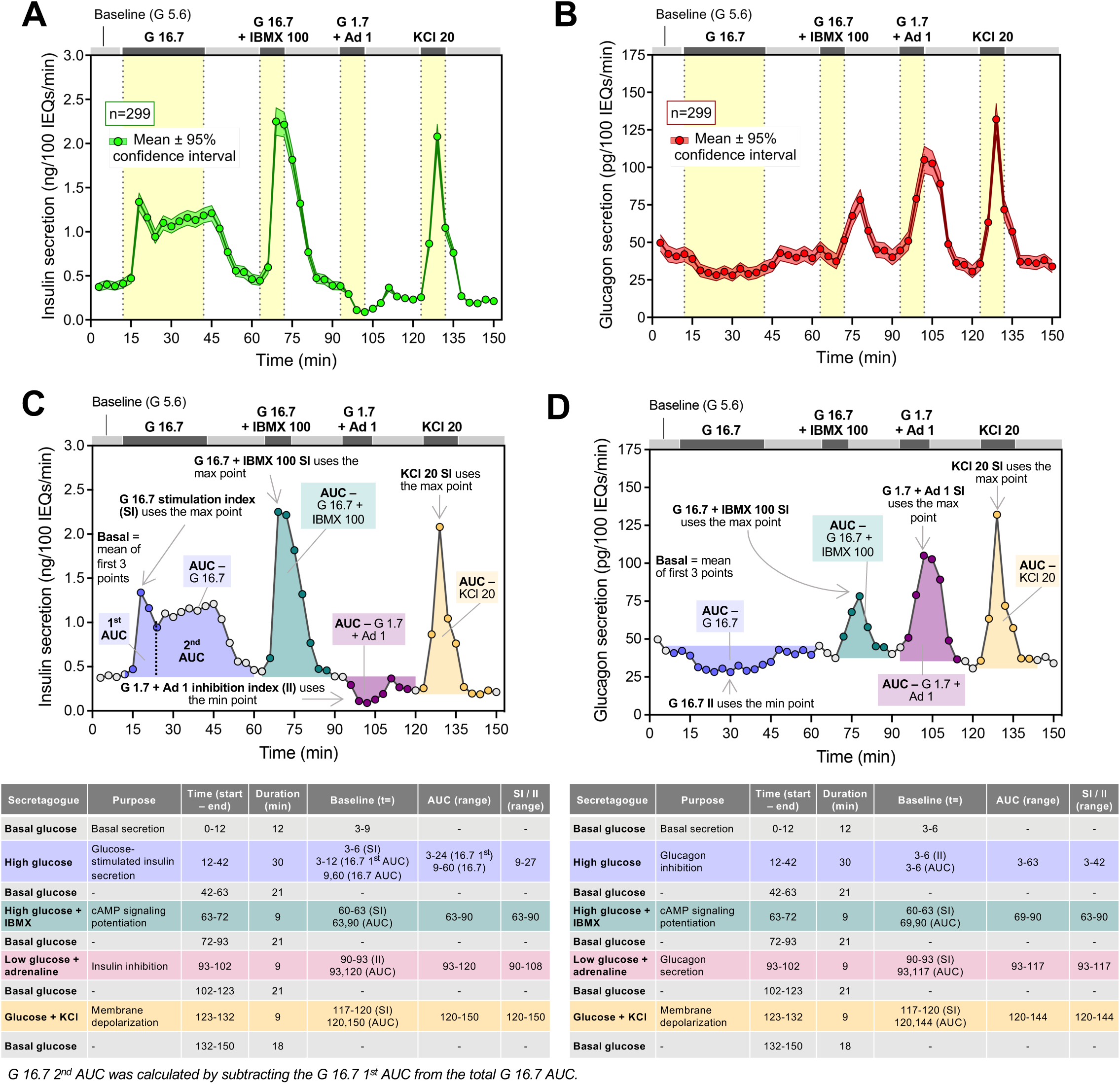
Islet secretory function is highly heterogeneous amongst donors. Average (**A**) insulin and (**B**) glucagon secretion from donor islets normalized to islet equivalents (IEQs). Data are displayed as mean ± 95% confidence interval (n = 299). (**C**) Schematic and tables describing insulin and (**D**) glucagon secretion traits derived from individual islet secretion traces. Descriptive statistics by sex and reported race/ethnicity are provided in **Extended Data Tables 1 and 2**. G – Glucose (mM); IBMX – isobutylmethylxanthine (μM); A – adrenaline (μM); KCl – potassium chloride (mM).

To understand how biological variables and islet processing features impacted hormone secretion across donors, we derived 11 insulin and 9 glucagon secretion traits from the respective hormone secretion traces (**Figure 2C-D**). For each human islet preparation, we derived the following secretion traits from each insulin trace: basal insulin secretion, glucose-stimulated 1^st^ phase secretion (G 16.7 1^st^ area under the curve; AUC), glucose-stimulated 2^nd^ phase secretion (G 16.7 2^nd^ phase secretion), overall glucose-stimulated insulin secretion (G 16.7 AUC), the glucose stimulation index (SI; G 16.7 SI), cAMP-potentiated insulin secretion (G 16.7 + IBMX 100 AUC), cAMP-potentiated SI (G 16.7 + IBMX 100 SI), response to low glucose plus adrenaline (G 1.7 + Ad 1 AUC), low glucose plus adrenaline inhibition index (II; G 1.7 + Ad 1 II), KCl-mediated insulin secretion (KCl 20 AUC), and KCl stimulation index (KCl 20 SI). We also derived the following secretory traits from each glucagon trace: basal secretion, glucose-inhibited glucagon secretion (G 16.7 AUC), glucose inhibition index (G 16.7 II), cAMP-potentiated glucagon secretion (G 16.7 + IBMX 100), cAMP-potentiated SI (G 16.7 + IBMX 100 SI), low glucose plus adrenaline-induced secretion (G 1.7 + Ad 1 AUC), low glucose plus adrenaline SI (G 1.7 + Ad 1 SI), KCl-mediated secretion (KCl 20 AUC), and the KCl stimulation index (KCl 20 SI). These data highlight the marked heterogeneity in responses, with the distribution of values shown for the total dataset and separately by sex and reported race and ethnicity or primary predicted ancestry in **Extended Data Tables 1-4**. Spearman correlation analyses revealed a significant correlation between donor age, sex, HbA1c, and BMI with multiple secretory traits (**Extended Data Figure 3**). Basal insulin secretion was positively correlated with HbA1c (r = 0.14, p_unadj_ = 0.02) and BMI (r = 0.14, p_unadj_ = 0.02). Age was negatively correlated with cAMP-potentiated insulin secretion (r = –0.15, p_unadj_ = 0.01) and the glucose stimulation index (r = –0.12, p_unadj_ = 0.047). The glucose stimulation index also was negatively correlated with sex (r = –0.12, p_unadj_ = 0.04), BMI (r = –0.14, p_unadj_ =0.02), and HbA1c (r = –0.14, p_unadj_ = 0.02). Additionally, HbA1c was positively correlated with cAMP-potentiated (r = 0.16, p_unadj_ = 0.005) and KCl-mediated glucagon secretion (r = 0.12, p_unadj_ = 0.04).

In multivariable models adjusting for donor age, sex, BMI, HbA1c, reported race or ethnicity, and islet isolation center, both pre-shipment culture time and islet transit time were significantly associated with multiple insulin and glucagon secretion traits (**Extended Data Figure 4A-B**). Given this, we added pre-shipment culture time and islet transit time into future multivariable regression models as additional covariates unless otherwise stated (8 covariates total). Of note, islet purity was associated with 3 islet functional traits; however, these data were unavailable for all observations for inclusion as a covariate in the regression model.

To remove potential confounding effects of other demographic and processing variables, we next explored associations between donor traits and secretory traits in multivariable analyses controlling for the other seven covariates. In these analyses, donor BMI was significantly associated with glucagon secretory traits (**Extended Data Figure 4D**). Specifically, BMI was negatively associated with basal (regression coefficient, b = –0.20, p = 0.005) and KCl-mediated depolarization of glucagon secretion (b = –0.17, p = 0.03) and positively associated with glucose-mediated glucagon inhibition (b = 0.15, p = 0.03; **Extended Data Figure 4D**). Donor age, sex, and reported race or ethnicity were associated with multiple secretory traits prior to, but not after adjusting for multiple comparisons (**Figure 3-4, Extended Data Figure 4E-F**). Interestingly, we found no differences in insulin or glucagon secretion between donors with HbA1c < 5.7% (n = 223) and those with HbA1c ≥ 5.7% (n = 76) (**Extended Data Figure 4C-D**).

**Figure 3.**
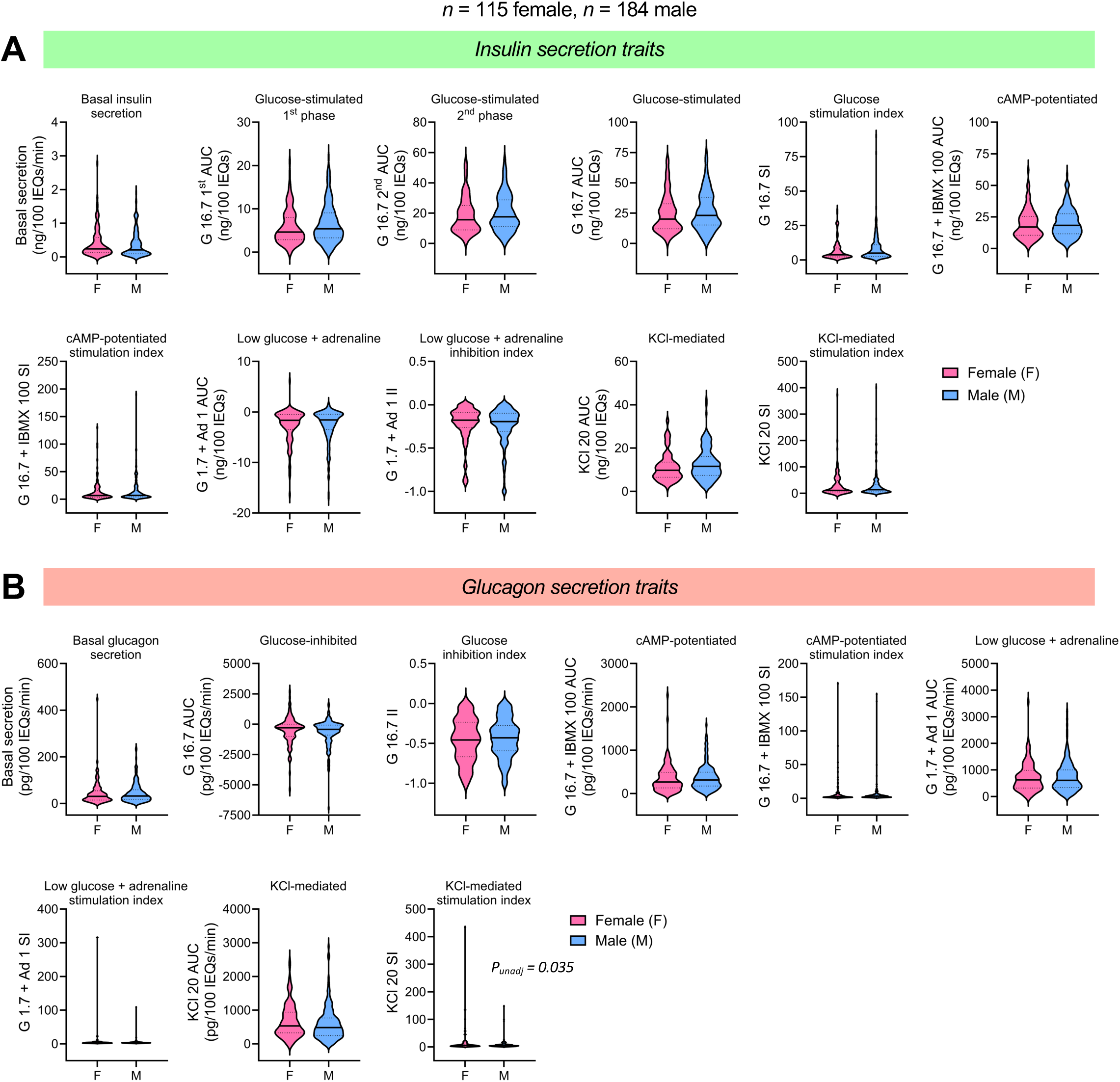
Islet secretion traits by sex. Violin plots comparing (**A**) insulin and (**B**) glucagon secretion traits between female (n = 115) and male (n = 184) donors. The solid line in each violin plot depicts the median, and the dotted lines represent the 1^st^ and 3^rd^ quartiles. Global P-value is based on the F-test while controlling for the seven covariates (age, sex, BMI, HbA1c, islet isolation center, pre-shipment culture time, and islet transit time) and adjusting for multiple comparisons. Comparisons where the unadjusted p-value (p_unadj_) was significant are noted in italics.

**Figure 4.**
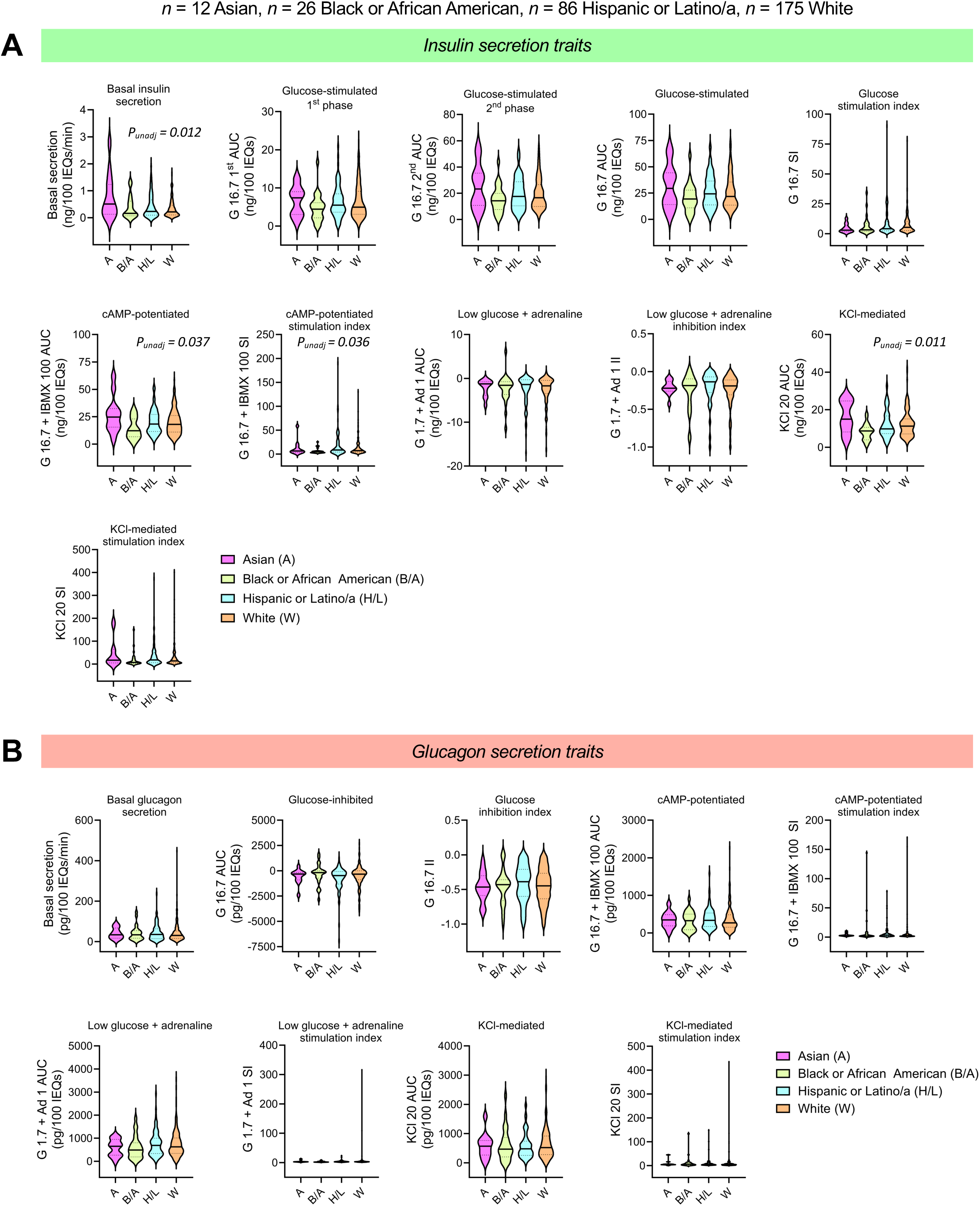
Islet secretion traits by reported race or ethnicity. Violin plots comparing (**A**) insulin and (**B**) glucagon secretion traits by reported race or ethnicity (n = 12 Asian, n = 26 Black or African American, n = 86 Hispanic or Latino/a, n = 175 White). The solid line in each violin plot depicts the median, and the dotted lines represent the 1^st^ and 3^rd^ quartiles. Global p-value is based on the F-test while controlling for the seven covariates (age, sex, BMI, HbA1c, islet isolation center, pre-shipment culture time, and islet transit time) and adjusting for multiple comparisons. Comparisons where the unadjusted p-value (p_unadj_) was significant are noted in italics.

For the human islet preparations for which genotyping data were available, we explored relationships between predicted genetic ancestry and islet function. We found that the relationships between primary predicted genetic ancestry and secretion were similar to those observed between reported race or ethnicity and insulin secretion. In addition, genetic ancestry had a statistically significant effect on basal insulin secretion (**Extended Data Figure 4E, Extended Data Figure 5A**).

### Islet composition strongly influences hormone secretory response

Next, we explored associations between islet function, morphology (i.e., diameter, area, perimeter), composition, and hormone content (**Figure 5**). Interestingly, in both Spearman correlation and multivariable analyses, islet composition was highly associated with multiple secretion traits, especially insulin secretion (**Figure 5A-B, Extended Data Figure 3**). A higher percentage of β cells was associated with an increase in glucose-stimulated (b = 0.23, p = 2.31 x 10^−4^), cAMP-potentiated (b = 0.35, p = 1.18 x 10^−8^), and depolarization-mediated insulin secretion (b = 0.30, p = 1.16 x 10^−6^). These secretory traits were negatively associated with the percentage of islet α and δ cells. Additionally, the percentage of δ cells within the islets was negatively associated with the glucose stimulation index (b = –0.13, p = 0.02) and inhibition index of insulin secretion in response to low glucose and adrenaline (b = –0.19, p = 1.76×10^−3^). These associations were marked by relatively large effect sizes, as evidenced by the absolute value of the scaled regression coefficient (0.13 – 0.35).

**Figure 5.**
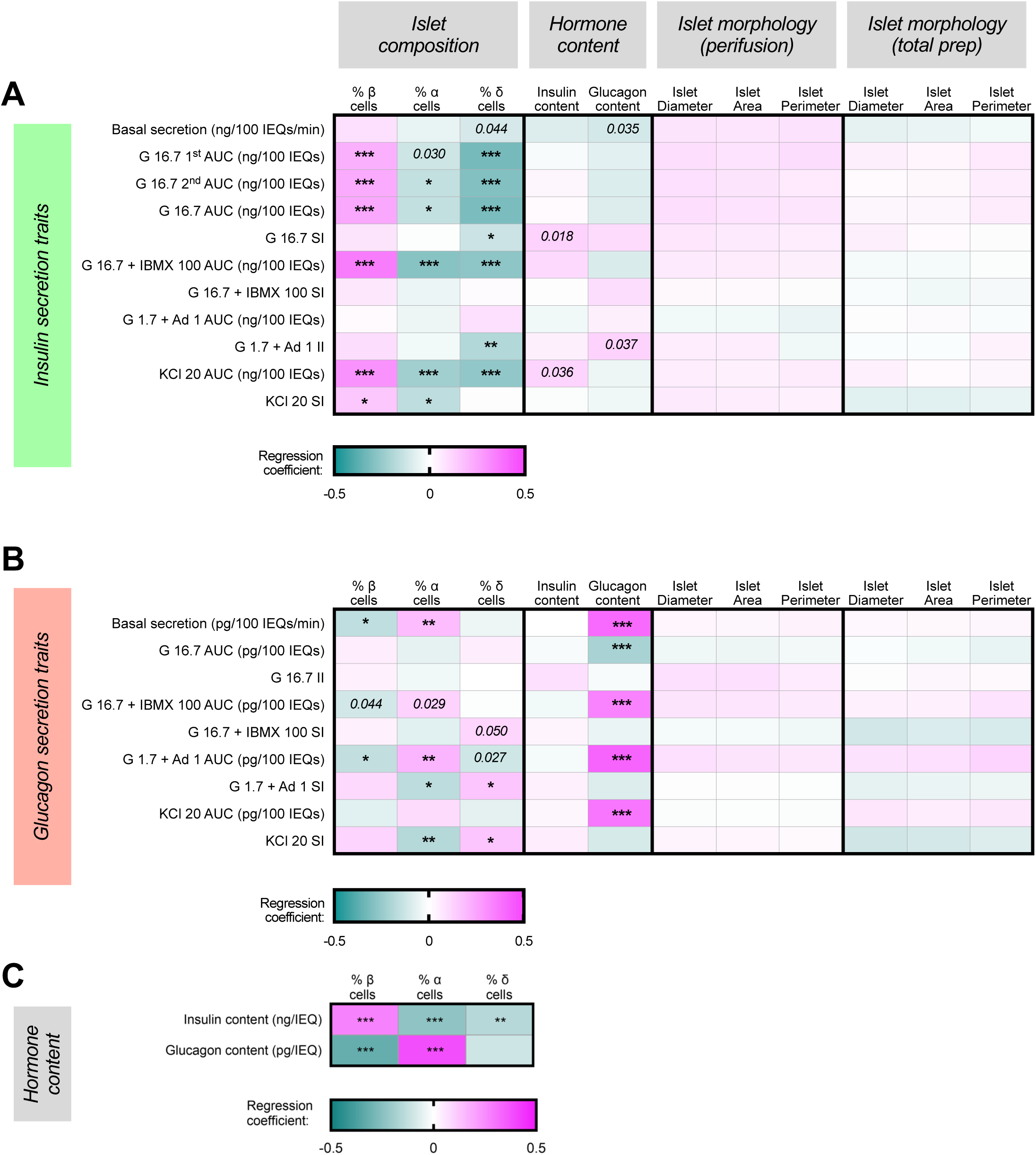
Islet composition and hormone content are associated with multiple islet secretory traits. (**A-B**) Heatmaps depicting regression coefficients of each morphologic variable for islets that underwent perifusion (n = 186), islet composition variable (n = 299), and total islet insulin and glucagon content (n = 299) after being incorporated into multivariable regression models controlling for age, sex, BMI, HbA1c, islet isolation center, reported race or ethnicity, pre-shipment culture time, and islet transit time for each (**A**) insulin or (**B**) glucagon secretion trait. (**C**) Heatmaps depicting regression coefficients of each islet composition variable after being incorporated into multivariable regression models for total insulin or glucagon content after adjusting for the eight covariates (n = 299). Adjusted p-values are indicated where *p < 0.05, **p < 0.01, and ***p < 0.001. Comparisons where the unadjusted p-value (p_unadj_) was significant are noted in italics.

Regarding the glucagon secretion traits, the percentage of α cells was positively associated with an increase in basal glucagon secretion (b = 0.18, p = 3.42 x10^−3^) and total glucagon secreted in response to low glucose with adrenaline (b = 0.21, p = 3.42 x10^−3^), while the percentage of β cells was negatively associated with these two traits (**Figure 5B**). Additionally, the percentage of α cells was negatively associated with the glucagon stimulation indices in response to both low glucose with adrenaline (b =-0.17, p = 1.58 x10^−2^) and KCl (b = – 0.18, p = 9.28 x10^−3^), while the opposite was true for their association with the percentage of δ cells.

Alternative analyses adjusting for population structure using the first five principal components explaining genetic ancestry as covariates, where available, yielded similar results (**Extended Data Figure 6**). Although a few associations were lost in these analyses, partially due to the reduced sample size, these analyses also revealed associations between the percentage of islet δ cells and the cAMP-potentiated glucagon stimulation index (b = 0.16, p = 2.05 x10^−2^) and the glucagon secreted in response to low glucose and epinephrine (b = –0.20, p = 8.31 x10^−3^) in **Extended Data Figure 6B**.

As expected, β and α cell composition were strongly associated with insulin and glucagon content, respectively (**Figure 5C, Extended Data Figure 3, Extended Data Figure 6C**). However, associations between hormone content were more striking for glucagon secretion traits than insulin secretion traits after controlling for the eight covariates and adjusting for multiple comparisons (**Figure 5A-B**). Glucagon content was strongly associated with multiple glucagon secretion traits (**Figure 5B, Extended Data Figure 6B**), including basal glucagon secretion (b = 0.40, p =1.13 x10^−13^) and total glucagon secreted in response to all four secretagogues; further, these associations were marked by relatively large effect sizes (0.21 – 0.42).

In contrast to the impact of islet composition and hormone content, we found no association between morphological traits (diameter, area, perimeter) of islets purified by hand-picking that underwent dynamic perifusion and any insulin or glucagon secretion traits (**Figure 5A-B**). Similar results were found when examining the relationship between the morphology-related traits of the islet preparation as a whole and islet functional traits (**Figure 5A-B**).

### Islet composition is influenced by sex, reported race/ethnicity, and predicted genetic ancestry

Given the strong association between islet composition and hormone secretion, we analyzed relationships between islet composition and donor characteristics. On average, analyzed islets were composed of 58% β cells (range 25 – 92%), 34% α cells (range 3 – 68%), and 8% δ cells (range 1 – 19%). Similar to islet hormone secretion, there was significant donor-to-donor heterogeneity in islet endocrine cell composition (**Figure 6A**). Donor sex and reported race or ethnicity, and genetic ancestry had a global effect on the percentage of islet β and α cells in multivariable models incorporating the other seven covariates (**Figure 6B-D**). Specifically, female sex was associated with a higher percentage of α cells and a lower percentage of β cells (**Figure 6B**). We observed a statistically significant global effect of reported race or ethnicity on the percentage of β cells (p = 0.003) and α cells (p = 0.003). Additonally, individuals reported as Asian in comparison to White (**Figure 6C**), as well as those with predicted East Asian in comparison to European predicted ancestry (**Figure 6D**), had a higher percentage of β cells (p = 0.0007 and p =0.0001, respectively) and a lower percentage of α cells (p = 0.0007 and p = 0.0001, respectively). There was a global effect of genetic ancestry on the percentage of β cells (p = 0.0004) and α cells (p = 0.0004) (**Extended Data Figure 7B**). We observed no significant effect of donor age, BMI, HbA1c, isolation center, pre-shipment culture time or islet transit time on islet composition in multivariable analyses (**Extended Data Figure 7A-B**).

**Figure 6.**
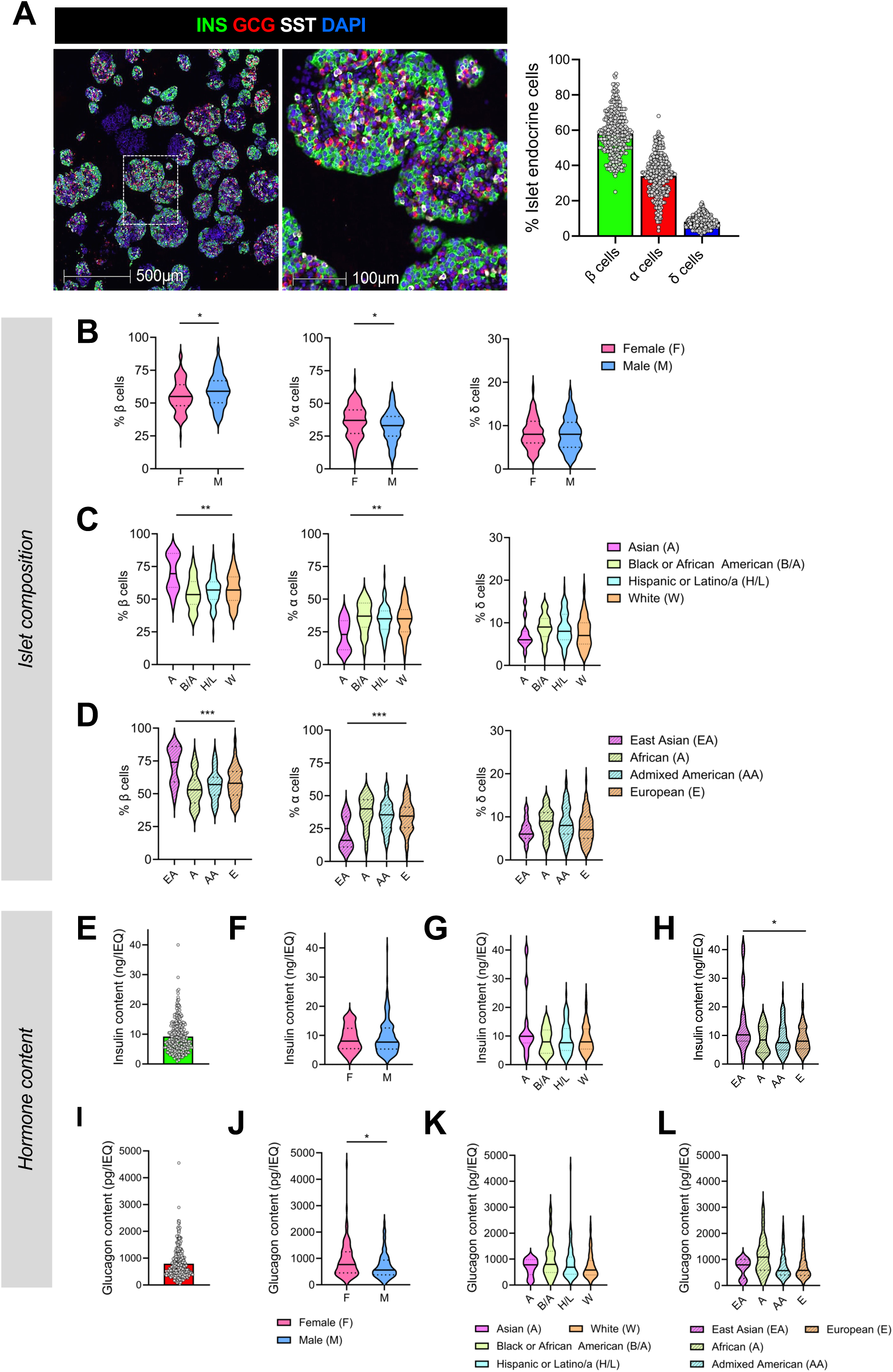
Islet composition and hormone content are highly variable and influenced by donor sex and ancestry. (**A**) Representative images of collagen gel-embedded islets and associated quantification of islet composition. Each point represents the average % β, % α, and % δ cells per donor (n = 299). Violin plots comparing islet composition by (**B**) donor sex, (**C**) reported race or ethnicity, and (**D**) predicted primary genetic ancestry. (**E**) Islet insulin content normalized to IEQ (n = 299). Violin plots comparing islet insulin content by (**F**) donor sex, (**G**) reported race or ethnicity, and (**H**) predicted primary genetic ancestry. (**F**) Islet glucagon content normalized to IEQ (n = 299). Violin plots comparing islet glucagon content by (**J**) donor sex, (**K**) reported race or ethnicity, and (**L**) predicted primary genetic ancestry. Donor sex: n = 115 female, n = 184 male. Reported race or ethnicity: n = 12 Asian, n = 26 Black or African American, n = 86 Hispanic or Latino/a, n = 175 White. Primary predicted genetic ancestry: n = 86 Admixed American, n = 21 African, n = 11 East Asian, n = 150 European. Adjusted global p-values are indicated where *p < 0.05, **p < 0.01, ***p < 0.001. For sex, reported race or ethnicity, and genetic ancestry, the global adjusted p-value is based on the F-test while controlling for the other seven covariates. INS – insulin; GCG – glucagon; SST – somatostatin.

We next investigated associations between islet hormone content, donor demographics, and islet processing traits (**Figure 6E-L, Extended Data Figure 7C-D**). We found a global effect of genetic ancestry on the insulin content (p = 0.0451). Additionally, individuals with predicted East Asian ancestry compared to those with European ancestry had higher insulin content (p = 0.0048) (**Figure 6H**). Insulin content was positively associated with donor age (b = 0.14, p = 0.0496) and negatively associated with transit time (b = –0.15, p = 0.03) in multivariable models controlling for the other seven covariates (**Extended Data Figure 7C-D**). Additionally, there was a global effect of genetic ancestry on insulin content (p = 0.045, **Extended Data Figure 7D**). In contrast, we found no associations between insulin content and donor sex, reported race or ethnicity, BMI, HbA1c, or isolation center in similar multivariable analyses (**Figure 6F-G; Extended Data Figure 7C-D**). Finally, female sex was associated with a higher glucagon content (**Figure 6J)** and there was a global effect of sex on glucagon content (p = 0.015; **Extended Data Figure 7D**), but no other associations were noted between glucagon content and other demographic or processing traits in multivariable models (**Figure 6K-L; Extended Data Figure 7C-D**). In analyses adjusting for population structure using the first five principal components explaining genetic ancestry, female sex was also associated with a slight increase in δ cell percentage (p = 0.049) (**Extended Data Figure 7E**), whereas there was no effect of sex on glucagon content (**Extended Data Figure 7F**).

### Genetic risk for diabetes is associated with islet composition

Both T1D and T2D GWAS signals map to loci associated with islet-enriched genes and their regulatory elements^6,37^. Furthermore, diabetes risk alleles have been associated with *in viv*o measures of islet function in participants without diabetes and linked to worsened islet function in those with T1D^38–40^. Thus, we were interested in determining how genetic risk for diabetes influenced *in vitro* hormone secretion, content, and islet composition in our study. In the current release of human islet preparations where donor genotyping data were available (n = 268), we generated T1D and T2D GRS, utilizing 67 and 338 single-nucleotide variants, respectively, based on previously published models^7,41^. Using these data, we investigated whether genetic risk for diabetes predicted any observed differences in islet function, composition, or hormone content in our dataset of human islet preparations from donors without diabetes. We further investigated the role of “process-specific” partitioned genetic risk scores previously published from clustering of intermediate traits to delineate heterogeneity in polygenic risk^42^. We incorporated either the complete or partitioned GRS into multivariable models that included the following as covariates with or without HbA1c: donor age, sex, BMI, HbA1c, isolation center, pre-shipment culture time, islet transit time, and the first five principal components explaining genetic ancestry (**Figure 7A-B; Extended Data Figure 8**). Interestingly, the T2D GRS was positively associated with the percentage of islet δ cells with HbA1c covariate (b = 0.17, p = 0.019; **Figure 7B**, *top*) or without HbA1c covariate (b = 0.17, p = 0.016; **Figure 7B**, *bottom*). At the same time, δ cells strongly influenced the islet hormone secretory traits (**Figure 5; Extended Data Figure 6**), especially those of β cells, suggesting that HbA1c does not mediate these associations. We detected no additional significant associations between T1D and T2D GRS and either hormone secretion or content when including HbA1c covariate in our analysis (**Extended Data Figure 8A-B**). In contrast, excluding HbA1c as a covariate not only detected a significant association between the T2D GRS and the percentage of islet δ cells (**Figure 7B***, bottom*) but also revealed a significant positive association between the T2D GRS and glucagon content (**Extended Data Figure 8D**). Individual data points for associations of % δ cells and T2D GRS are shown in **Extended Data Figure 9A**.

**Figure 7.**
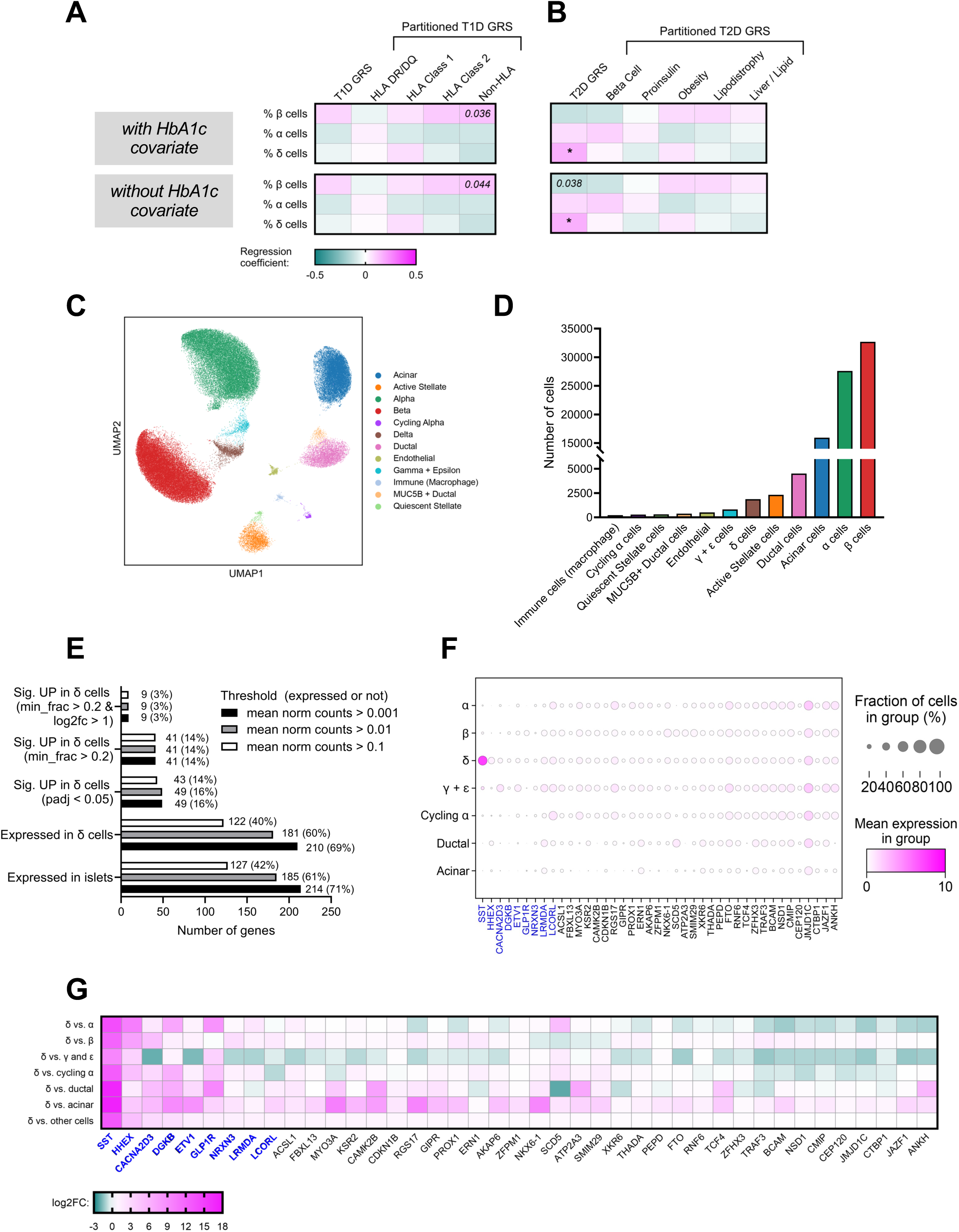
Pancreatic islet δ cell composition is associated with a genetic risk score for type 2 diabetes derived from genome-wide association studies. (**A-B**) Heatmaps depicting coefficients of the GRS for (**A**) T1D and (**B**) T2D, along with partitioned GRS variables after being incorporated into multivariable regression models for each secretion trait, hormone content, and islet composition variable (n = 268). Each model included the following covariates:donor age, sex, BMI, islet isolation center, pre-shipment culture time, islet transit time, and the first five principal components explaining primary genetic ancestry, with the HbA1c covariate (*top*) or without HbA1c covariate (*bottom*). “T1D GRS” encompasses the genetic risk for T1D (67 total variants). “HLA DR/DQ” represents the sum of T1D risk at HLA-DR/DQ haplotypes. “HLA Class 1” and “HLA Class 2” represent the sum of T1D risk at HLA class 1 and 2 alleles, respectively, excluding HLA DR/DQ haplotypes. “Non-HLA” represents the sum of T1D risk from variants outside the HLA region, genome-wide. “T2D GRS” encompasses genetic risk for T2D (338 total variants). “Beta Cell,” “Proinsulin,” “Obesity,” “Lipodystrophy,” and “Liver/Lipid” represent partitioned T2D GRS clusters encoding the related variants. Adjusted p-values are indicated where *p < 0.05. Comparisons where the unadjusted p-value (p_unadj_) was significant are noted in italics. (**C**) Clustering of the HPAP single-cell RNA-seq data from 44 HPAP donors without diabetes. (**D**) The number of cells of each type included in the analysis, including 1877 δ cells. (**E**) The number of genes that were differentially expressed, with three different thresholds for expression, in δ cells. (**F**) A dot plot showing T2D GRS genes that are differentially expressed in δ cells (significant genes are shown in blue). (**G**) A heat map illustrating T2D GRS genes that are differentially expressed in δ cells (significant genes are shown in blue).

To further establish whether the underlying transcripts at T2D GWAS signals captured within the T2D GRS were enriched in δ cells, we analyzed 87,505 islet cells (1877 δ cells) from 44 donors without diabetes using single-cell RNA sequencing data from the Human Pancreas Analysis Program (HPAP)^43–46^ (**Figure 7C-G**). Since the precise effector transcript remains unknown for many T2D GWAS signals, we used the proximal gene to evaluate the polygenic contributions of the 338 GRS variants (**Figure 7E** and **Extended Data Figure 9B**). Our analysis demonstrated that several T2D GRS candidate genes are strongly upregulated in δ cells (**Figure 7F**, significant genes highlighted in blue). Furthermore, we showed that the T2D GRS is highly enriched for genes that are strongly expressed in δ cells (**Figure 7G**, with significant genes highlighted in blue, and **Extended Data Figure 9C**).

## DISCUSSION

Islet physiology is disrupted in both T1D and T2D. The pathways and cell-cell communications that underlie islet dysfunction in diabetes are poorly understood, yet they may have important implications in treatment strategies employing both pharmacologic agents and islet or β cell replacement therapy. Recent emphasis on precision medicine approaches coupled with increased utilization of single-cell technologies has highlighted an important role for heterogeneity in diabetes classification, treatment, and molecular phenotypes^47–51^. To understand how biological variation in humans impacts islet function and diabetes risk, we generated this research resource from islets distributed through the NIH-funded IIDP. These islets underwent pre-shipment assessment at one of five islet isolation centers followed by centralized phenotyping and genotyping through the HIPP and HIGI programs. A strength of our study is the inclusion of islets from a diverse group of U.S. organ donors with over 40% of donors having a reported race or ethnicity other than White, non-Hispanic or Latino/a. The IIDP-HIPP-HIGI workflow has enabled investigation for associations between inherent donor characteristics, islet composition, genetic risk scores for T1D and T2D, and multiple insulin and glucagon secretory traits. Our data can be accessed through the IIDP RDR, where researchers can view and interrogate the dataset used for this analysis and propose their own unique questions related to human islet biology using the larger RDR dataset. Images presented in this report are available on the open-access Pancreatlas™ platform^52^, which supports interactive exploration of full-resolution islet imaging data.

Our phenotyping workflow includes an assessment of islet purity and viability, insulin and glucagon secretion measured in a dynamic perifusion system, and a quantitative assessment of islet composition in a sample of islets distinct from the islet aliquot used for perifusion. Studies of the human pancreas suggest a greater than four-fold variation in β cell mass amongst individuals without diabetes^53^. Not surprisingly, we observed a wide range of β cell secretory responses to glucose, IBMX, and KCl, in line with previous reports ^54–56^. While clinical studies suggest that glucose tolerance declines with age, whether β cell function declines with age independently of changes in insulin resistance and weight in individuals who maintain normal glucose tolerance is not clear^57,58^. Similarly, the impact of donor age on insulin secretion *in vitro* has been mixed in previous reports, with studies finding either no effect or a negative impact of age on glucose-stimulated insulin secretion^55,59–62^. In addition, many studies report an effect of BMI on multiple insulin secretory traits, while others have found no correlation^54,55,61,63^. In the current study, Spearman correlation analyses revealed a significant positive relationship between BMI and basal insulin secretion and a negative relationship between either BMI or age and the glucose stimulation index. However, we detected no significant association between any donor trait and insulin secretion in multivariable regression models adjusting for 7 other donor and islet processing-related covariates. Our studies differ from these previous studies in that we included a larger sample size of donors and controlled for multiple potentially confounding demographic and islet processing features, which we and others have shown to impact *in vitro* secretory responses^55,61,64^.

In contrast to the weak effects of donor demographics on insulin secretion, islet composition exhibited strong and consistent associations with insulin secretory traits in both Spearman correlation and multivariable analyses. Consistent with observed heterogeneity in hormone secretory responses, islet composition was highly heterogeneous amongst islet preparations, with an approximately 4, 22, and 19-fold variation in the percentages of β, α, and δ cells, respectively. β cell composition or a higher percentage of β cells was associated with increased glucose-stimulated, cAMP-potentiated, and depolarization-mediated insulin secretion. While islet isolation severs vascular and neuronal connections, the autonomy of the islet as a mini-organ is well preserved after isolation, as demonstrated by the persistence of paracrine interactions and electrical coupling that modulate islet cell-cell communication within islets^65^.

Strikingly, although δ cells comprised a relatively low percentage of islet endocrine cell composition, a higher percentage of δ cells negatively impacted most insulin secretion traits. This finding is notable given a recent report showing that δ cells regulate the glycemic set point in healthy mice and that ablation of the δ cell population leads to blood glucose lowering and increased insulin secretion^66^. In multivariate analysis, we queried factors associated with T1D and T2D GRS, identifying a positive association between the percentage of islet δ cells and the T2D GRS. Of the 303 candidate genes that map to the 338 variants reflected in the T2D GRS, 181 of 303 (60%) genes were expressed (mean of normalized counts > 0.01) in δ cells. Notable genes identified in this analysis include the transcription factor *HHEX*, which has previously been shown to play an essential role in δ cell differentiation, with loss of *HHEX* leading to a reduced number of δ cells, reduced islet somatostatin secretion, and increased insulin secretion^67^. We identified δ cell expression of other notable genes with resolved effector transcript status, including *DGKB*^4^ and *GLP1R*^68,69^, which have been associated with glycemia and insulin secretory phenotypes in human studies.

Multiple glucagon secretory traits were associated with islet composition; however, we found even stronger associations between glucagon content and total glucagon secreted in response to all secretagogues used in this study. Interestingly, this same relationship was not observed between insulin content and insulin secretory responses. One study comparing murine β and α cells noted faster rates of glucagon granule exocytosis, larger readily releasable pools, and a higher rate of refilling of these pools after depletion in α cells^70^. Therefore, it is possible that these same mechanisms are at play in human islets and could explain differences in the association between hormone content and secretion between β and α cells.

Our observations on the impact of islet composition on secretory traits raise an important question of whether differences in cell composition are influenced by donor demographics and/or genetics. Interestingly, we found a global effect of sex and reported race or ethnicity on the percentage of islet β and α cells. Here, female donors had a higher percentage of α cells, and donors reported as Asian exhibited a higher percentage of β cells. We note the relatively small number of donors reported as Asian in the dataset, highlighting a need to confirm this observation with larger sample sizes. Remarkably, a high T2D GRS was positively associated with the percentage of islet δ cells, which was consistent with the finding that δ cell composition negatively impacted most insulin secretion traits, highlighting the inhibitory potential of δ cell-secreted factors, including somatostatin. Together, these data suggest that, in some individuals, a genetic predisposition to having more δ cells may impact insulin secretory capacity and glycemic setpoints, making one more susceptible to the development of T2D. This is further supported by the structural basis for δ cell paracrine regulation in pancreatic islets^71^. However, the mechanisms by which genetic variation associated with altered T2D risk influences δ cell composition are unknown. Recent studies have utilized the abundance of known endocrine marker proteins to infer associations with gene and protein expression^56,64^; however, this is the first time to our knowledge that direct quantitative assessment of islet composition has been compared to hormone secretion.

In summary, this study has identified important relationships between donor demographics, genetics, islet processing, islet composition, hormone content, function, using human islet preparations from donors without diabetes. Given their impact on human islet function, these factors should be considered in interpreting future human islet studies. Further, they may have implications in strategies for β cell replacement therapy^72–74^. For example, in 2022 the FDA approved Lantidra^75^, the first allogeneic pancreatic islet cellular therapy made from deceased donor pancreatic cells for the treatment of T1D to ameliorate problematic hypoglycemia with glucocorticoid-free immunosuppression^76^. Release criteria for islet grafts have historically included measures of glucose-stimulated insulin secretion, which predicts metabolic outcomes of human islet transplantation^77^. Our data highlight the impact of cell-cell interactions on islet function and suggest that islet composition may provide valuable insight into graft function post-transplant. Similarly, we expect that integrating these findings from 299 donors without diabetes with datasets generated using the same phenotyping and genotyping pipelines for islets from individuals with T1D and T2D, such as those generated by the Human Pancreas Analysis Program (HPAP)^43,46^ will inform our understanding of diabetes pathophysiology and lead to the design of new therapies and disease prevention.

There are limitations of our analysis that should be acknowledged. First, our analysis was performed on islets isolated from the pancreas of organ donors following brain or circulatory death. Because the pancreas cannot be safely biopsied in living individuals, the analysis of post-mortem tissue remains an important caveat for most human islet research. While we evaluated the impact of donor demographic features and technical features of the isolation, we did not evaluate the impact of the reported cause of death on islet function or viability. There is some data to suggest that islet secretory function varies by category of reported cause of death (i.e., trauma, cerebrovascular, or cardiovascular)^78^. However, autopsy studies indicate that clinical assessment of the cause of death is prone to error in an estimated 13-38% of cases^79^. Further, whilst we examined factors impacting glucose-stimulated islet responses in detail, we did not investigate responses to other nutrients, such as fatty acids or amino acids, which also display significant donor-to-donor heterogeneity, as highlighted by a recent report from Kolic et al.^56^. Finally, while this study is currently the largest of its type, we had multiple observations that were nominally significant, supporting the need for larger studies and expanded sample sizes. The diverse IIDP-HIPP-HIGI datasets represent a valuable and continuously growing resource for the islet biology community, which can be used to generate new scientific hypotheses and guide mechanistic studies of human islets. Future studies should leverage this rich dataset to link islet secretory characteristics and composition with multi-omic assessment and test these relationships in cohorts of donors, including controls and those with pre-diabetes and diabetes, especially to understand cell-specific contributions to diabetes risk and disease pathophysiology.

## RESOURCE AVAILABILITY

### Lead contact

Further information and requests for resources and reagents should be directed to and will be fulfilled by the lead contact, Marcela Brissova, PhD. Requests for access to genetic data should be directed to Anna L. Gloyn, DPhil.

### Materials availability

This study did not generate new, unique reagents.

### Data and code availability

Data from the first release of 299 de-identified human islet preparations reported in this paper is available in **Supplemental File 2.** The genetic risk scores and predicted ancestry for each donor and the larger integrated dataset hosted are available for download from the IIDP Research Data Repository. For more information on available datasets and how to access, visit https://iidp.coh.org/Resources-Offered/Research-Data-Repository. Genotyping data were deposited in the European Genome Phenome Archive (EGA) under accession number EGAS50000000697 and are available through a data access agreement. All original code is publicly available as of the date of publication. Code related to the generation of statistical models is available at https://github.com/hakmook/HIPP_project. Code associated with the prediction of genetic ancestry is available at https://github.com/gloynlab/GeneticAncestry. Any additional information required to reanalyze the data reported in this paper is available from either Dr. Brissova or Dr. Gloyn (genetic data) upon request.

## Supporting information

Extanded Data Figures and Tables

## ACKNOWLEDGMENTS

The authors would like to thank the organ donors and their families for their invaluable donations and acknowledge the IIDP collaborators who continue to make this work possible (**Supplemental File 1**). The authors would also like to thank Danielle Gibson (Vanderbilt University Medical Center) for her assistance with the HIPP studies, Dr. Emily Anderson-Baucum (Indiana University School of Medicine) for her helpful advice and edits, and Drs. Ha Vu, Stephen Parker, and Jie Liu for their critical reading of the manuscript and contributions to the edits. Research reported in this publication was supported by U42RR017673, UC4DK098085, U24DK098085, 1-RSC-2018-561-I-X, 1-RSC-2019-712-S-B, 1-RSC-2020-890-S-B, F30DK134041, T32GM007347, T32GM152284, R01DK129469, and The Leona M. and Harry B. Helmsley Charitable Trust. Whole-slide imaging was performed in the Islet and Pancreas Analysis Core of the Vanderbilt DRTC (DK20593).

We also acknowledge the resources provided by PanKbase (https://www.pankbase.org), which is supported by NIDDK U24DK138515 and U24DK138512, as well as supplemental funds from the NIH Office of Data Science Strategy (U01DK128847). This manuscript used data acquired from the database (https://hpap.pmacs.upenn.edu/) of the Human Pancreas Analysis Program (HPAP; RRID:SCR_016202). HPAP is part of a Human Islet Research Network (RRID:SCR_014393) consortium (UC4DK112217, U01DK123594, UC4DK112232, and U01DK123716).

## AUTHOR CONTRIBUTIONS

Conceptualization: M.B., Y.D.P., H.K., D.C.S., S.A.S., A.L.G., J.C.N., and C.E.M.; Data curation: Y.D.P., D.C.S., S.A.S., H.S., A.L.H., A.B., K.X., and F.F.; Formal analysis: H.K., D.S., K.X., Y.D.P., S.A.S., H.S., and D.C.S.; Funding acquisition: C.E.M., K.A., A.L.G., J.C.N., and M.B.; Investigation: H.D., S.M., A.C., C.D., C.V.R., J.T., T.SR.B., R.A., G.P., A.E., R.J. V.S., and S.T.; Methodology: S.A.S., H.S., D.S., H.K., A.C.P., S.A.S. H.S. A.L.G and M.B.; Project administration: C.E.M., K.A., A.L.G., J.C.N., and M.B.; Software: JP.C.; Resources: M.B., C.E.M., A.L.G., J.C.N., JP.C., and A.C.P.; Supervision: M.B., C.E.M., A.L.G, J.C.N., H.K., A.C.P., and K.A.; Visualization: Y.D.P., S.A.S., H.S., and D.C.S.; Writing – original draft: Y.D.P., C.E.M., A.L.G., and M.B.; Writing – review & editing: all authors.

## EXTENDED DATA FIGURE AND TABLE LEGENDS

**Extended Data Figure 1, related to Figure 1**. Schematic of multimodal data integration by phenotyping and genotyping pipelines within Integrated Islet Distribution Program (IIDP) and data deposition in Resource Data Repository (RDR).

**Extended Data Figure 2, related to Figure 1. Example HIPP procedures.** (**A**) Representative images of a human islet preparation stained with dithizone (DTZ) to delineate islet versus non-islet tissue. Using the Count and Measure function in cellSens, images of DTZ-stained islets were utilized to quantify the total islet equivalents (IEQs), purity, and islet morphology. Tissue depicted in red on the image mark-up indicates positively identified islet tissue, while green marks non-islet tissue. (**B**) Representative images of handpicked islets pre– and post-perifusion. The red color on the image markup indicated positively identified islet tissue by the Count and Measure function in cellSens. The scale bar is 500 μm.

**Extended Data Table 1, related to Figure 2. Insulin secretion is highly heterogeneous amongst donors.** Descriptive statistics of the eleven insulin secretion traits by donor sex and reported race or ethnicity. Data are displayed as mean ± SD, followed by the range.

**Extended Data Table 2, related to Figure 2. Glucagon secretion is highly heterogeneous amongst donors.** Descriptive statistics of the nine glucagon secretion traits by donor sex and reported race or ethnicity. Data are displayed as mean ± SD, followed by the range.

**Extended Data Table 3, related to Figure 2. Distribution of insulin secretion traits is similar in n = 268 subset.** Descriptive statistics of the eleven insulin secretion traits by donor sex and genetic ancestry. Data are displayed as mean ± SD, followed by the range.

**Extended Data Table 4, related to Figure 2. Distribution of glucagon secretion traits is similar in n = 268 subset.** Descriptive statistics of the nine glucagon secretion traits by donor sex and genetic ancestry. Data are displayed as mean ± SD, followed by the range.

**Extended Data Figure 3, related to Figure 2. Relationship between donor demographics, islet processing, islet morphology, and functional traits.** Heatmap of Spearman r correlation coefficients between the indicated variables in the corresponding labeled row and column. For donor sex, the male was coded as 0 and female as 1. *p < 0.05; †p < 0.01; ‡p < 0.001. All p-values are unadjusted.

**Extended Data Figure 4, related to Figures 3 and 4. Donor and islet processing traits correlate with islet function.** (**A-B**) Heatmaps depicting regression coefficients of each islet processing trait after being incorporated into multivariable regression models for each (**A**) insulin and (**B**) glucagon secretion trait. Each model included the following covariates: age, sex, reported race or ethnicity, BMI, HbA1c, and islet isolation center. Pre-shipment culture time and islet transit time were included as additional covariates for all subsequent models, unless noted otherwise. (**C-D**) Heatmaps depicting regression coefficients of donor age, BMI, and prediabetes status after being incorporated into multivariable regression models for each (**C**) insulin and (**D**) glucagon secretion trait while controlling for the other seven final covariates. Adjusted p-values are indicated where *p < 0.05, **p < 0.01, and **p < 0.001. For associations that are significant prior to adjusting to multiple comparisons, the unadjusted p-value is noted in italics. (**E-F**) Tables summarizing the global adjusted P-value when comparing differences by donor sex, reported race or ethnicity, genetic ancestry, and isolation center for each (**E**) insulin and (**F**) glucagon secretion trait. The global adjusted p-value is based on the F-test while controlling for the other seven covariates. Analyses involving genetic ancestry controlled for all covariates except reported race or ethnicity. n.s. indicates an insignificant global adjusted p-value and unadjusted p-value. For results that were significant prior to adjusting to multiple comparisons, the unadjusted p-value is noted in italics. n = 299 for all analyses except those involving islet purity (n = 269), elevated HbA1c (n = 76 elevated HbA1c, n = 223 normal HbA1c), and genetic ancestry (n = 268).

**Extended Data Figure 5, related to Figure 4. Islet secretion traits by genetic ancestry.** Violin plots comparing (**A**) insulin and (**B**) glucagon secretion traits by primary predicted genetic ancestry (n = 86 Admixed American, n = 21 African, n = 11 East Asian, n =150 European). The solid line in each violin plot depicts the median, and the dotted lines represent the 1^st^ and 3^rd^ quartiles. *p < 0.05 after adjusting for multiple comparisons. Comparisons where the unadjusted p-value (p_unadj_) was significant are noted in italics. Global p-value is based on the F-test while controlling for the seven covariates (donor age, sex, BMI, HbA1c, islet isolation center, pre-shipment culture time, and islet transit time).

**Extended Data Figure 6, Related to Figure 5. Islet composition and hormone content are associated with multiple islet secretory traits after adjusting for population structure using the first five principal components explaining genetic ancestry.** (**A-B**) Heatmaps depicting regression coefficients of each morphologic variable for islets that underwent perifusion (n = 186), islet composition variable (n = 268), and total islet insulin and glucagon content (n = 268) after being incorporated into multivariable regression models controlling for age, sex, BMI, HbA1C, islet isolation center, islet transit time, pre-shipment culture time and the first 5 principial components explaining genetic ancestry for each (**A**) insulin or (**B**) glucagon secretion trait. (**C**) Heatmaps depicting regression coefficients of each islet composition variable after being incorporated into multivariable regression models for total insulin or glucagon content after adjusting for the eight covariates (n = 268). Adjusted p-values are indicated where *p < 0.05, **p < 0.01, and ***p < 0.001. Comparisons where the unadjusted p-value (p_unadj_) was significant are noted in italics.

**Extended Data Figure 7, related to Figure 6. Correlation between donor demographics and islet processing variables with islet composition and hormone content.** (**A**) Heatmaps depicting regression coefficients of donor and islet processing variables when incorporated into multivariable regression models with an islet composition variable (% β, α, or δ cells) as the outcome variable. For models related to islet processing traits, the following covariates were included: donor age, sex, reported race or ethnicity, BMI, HbA1c, and islet isolation center. For donor traits, pre-shipment culture time and islet transit time were also included as covariates. For associations that are significant before adjusting for multiple comparisons, the unadjusted p-value is noted in italics. (**B**) Table summarizing the global adjusted p-value when comparing differences by isolation center for each composition variable. The global adjusted p-value is based on the F-test while controlling for the other seven covariates listed in **A**. (**C**) Heatmaps depicting regression coefficients of donor and islet processing variables when incorporating into multivariable regression models, insulin and glucagon content. For models related to islet processing traits, the following covariates were included: age, sex, reported race or ethnicity, BMI, HbA1c, and islet isolation center. For donor traits, pre-shipment culture time and islet transit time were also included as covariates. Adjusted p-values are indicated where *p < 0.05. (**D**) Table summarizing the global adjusted p-value when comparing differences by isolation center for hormone content. The global adjusted p-value is based on the F-test while controlling for the other seven covariates listed in **C**. (**E-F**) Violin plots comparing islet composition (**E**) and glucagon content (**F**) by donor sex when adjusted for population structure, including seven covariates: age, sex, BMI, HbA1c, islet isolation center, islet transit time, pre-shipment culture time, plus 1^st^ five principal components explaining genetic ancestry. Global adjusted p-values based on the F-test while controlling for the other twelve covariates are indicated where *p < 0.05. For analyses including genetic ancestry as a covariate, n = 268; otherwise, n = 299 for all other analyses.

**Extended Data Figure 8, related to Figure 7. Associations between T1D and T2D GRSs and islet function and hormone content, including and disregarding HbA1c. (A-B)** Heatmaps depicting coefficients of the GRS for (**A**) T1D and (**B**) T2D along with partitioned GRS variables after being incorporated into multivariable regression models for each secretion, hormone content, and islet composition variable (n = 268). Each model included the following covariates: donor age, sex, BMI, islet isolation center, pre-shipment culture time, islet transit time, and the first five principal components explaining primary genetic ancestry. Adjusted p-values are indicated where *p < 0.05. For associations that are significant before changing to multiple comparisons, the unadjusted p-value is noted in italics.

**Extended Data Figure 9, related to Figure 7. Mapping predicted T2D GWAS effector transcripts in islet δ cells.** (**A**) Visualization of % δ cell composition in islets vs. T2D GRS. The data were used for associations of % δ cells and T2D GRS after adjusting for the following covariates: donor age, sex, BMI, islet isolation center, pre-shipment culture time, islet transit time, and the first five principal components explaining primary genetic ancestry, with or without HbA1c. (**B**) Number of signals included in the T2D GRS that map to an individual gene. Some genes have more than one signal; therefore, 338 variants were mapped to 303 genes. (**C**) Enrichment of T2D GRS for genes strongly expressed in δ cells.

## METHODS

### EXPERIMENTAL MODEL AND STUDY PARTICIPANT DETAILS

#### Primary human islet isolation

Human islet preparations from cadaveric organ donors (n = 299) were isolated from five affiliated islet isolation centers across the U.S and distributed to the Human Islet Phenotyping Program through the IIDP from 2016-2024. Donor demographic information is detailed in **Figure 1**. De-identified human pancreatic specimens do not qualify as human subjects research under the Vanderbilt or the Stanford Institutional Review Boards. All assays were performed on the day of islet arrival. Islet preparations included in this manuscript (n = 299) were isolated from five IIDP-affiliated isolation centers: Scharp-Lacy (n = 113), Southern California Islet Cell Resource Center (n = 94), University of Miami (n = 30), University of Pennsylvania Islet Transplant Center (n = 27), and the University of Wisconsin Human Islet Core (n = 35).

### METHOD DETAILS

All human islet preparations were assessed according to standardized protocols established by the HIPP and HIGI of the IIDP and are publicly available at https://iidp.coh.org/SOPs. All HIPP-related assessments are conducted on the day of arrival. Methods are briefly summarized below.

#### Islet purity and morphology

A representative sample of the unpicked human islet preparation was prepared by combining a 320 μL aliquot of islet preparation and 680 μL CMRL medium. The representative sample was stained with 50 μL of 1 mg/mL dithizone/DMSO/PBS (DTZ; Sigma, catalog #D5130) for 1-2 min at room temperature. After adding 1 mL of CMRL medium, brightfield and darkfield images of stained islets were captured at 10X magnification. Islet (stained) and non-islet (unstained) area was determined using the Count and Measure function in cellSens (Olympus). Islet purity was calculated as the following:

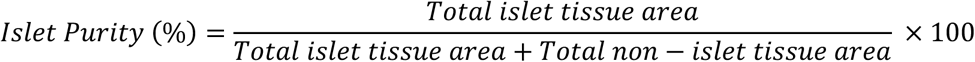

Using the Count and Measure function in cellSens, morphological characteristics of each islet were also determined, including average islet radius, diameter, perimeter, and area.

Measurements were performed in triplicate to ensure accuracy.

#### Islet viability

Qualitative assessment of islet viability upon arrival to the HIPP was performed using fluorescein diacetate (FDA) and propidium iodide (PI). In brief, 400 µL of well-mixed total islet suspension was added to a cell culture dish, and 10 µL of PI (Sigma Aldrich, catalog # P4170) and 10 µL of FDA (Sigma Aldrich, catalog # F-7378) were added to the islet suspension in succession. The plate was incubated in the dark for 15 mins, and the preparation was imaged immediately using an Olympus SZX12 stereomicroscope system. Multiple fields of view were captured from two replicates per donor to ensure that FDA/PI staining was visualized in 50-100 islets per preparation.

Additionally, a quantitative assessment of islet viability upon arrival to the HIPP was performed using trypan blue dye. Two aliquots of ∼100 handpicked islets were prepared in 1.5 mL tubes and washed three times using 2 mM EDTA/PBS via centrifugation at 200 rcf for 1 min at 4 °C. Islets were dispersed into a single cell suspension using Accutase (Innovative Cell Technologies, Inc, catalog # AT-104) and triturated at a slow speed for 10 minutes using an electronic multichannel pipet with tips. The reaction was quenched with CMRL-1066 media containing 10% FBS, 2 mM L-glutamine, and 1% penicillin/streptomycin, and cells were washed twice via centrifugation at 500 rcf for 3 mins at 4 °C. The cell pellet was resuspended in 50-75 µL of CMRL-1066 medium by gently flicking the tube. Next, 10 μL of the cell suspension was transferred to a new 1.5 mL tube and mixed with 10 µL Trypan Blue dye (Invitrogen, catalog # T10282). The cell chamber slide was loaded with 10 µL of the mixture in each compartment, and cell counts were acquired using the Countess II cell counter following instrument guidelines. Cell counts were repeated for the second aliquot of handpicked islets. Reported viability represents the average percentage of viable cells for duplicates of purified islets based on two cell counts per aliquot.

#### Islet function in a dynamic perifusion system

Islet function was assessed via dynamic perifusion. On the day islets were received, half of the islet shipment (1000-5000 islet equivalents (IEQs)) was plated in a 10-cm non-tissue culture treated dish and cultured in CMRL-1066 plus 10% FBS media at 37 °C and 5% CO_2_ for 2 hours prior to perifusion. Under microscope guidance, 267-300 IEQs were handpicked, islet size and count were recorded, and high-resolution brightfield and darkfield images were captured at 10x magnification. All islets were then transferred to a clear Eppendorf tube for loading into the perifusion chamber, fractions were collected at 1 mL/min, and secretagogues were changed at predetermined fractions. The following secretagogues were used for these perifusion studies: 5.6 mM glucose (baseline media; G 5.6), 16.7 mM glucose (G 16.7), 16.7 mM glucose with 100 μM isobutylmethylxanthine (IBMX; G 16.7 + IBMX 100), 1.7 mM glucose with 1 μM adrenaline (G 1.7 + Ad 1), and 5.6 mM glucose with 20 mM KCl (G 5.6 + KCl 20). All secretagogues were prepared in a perifusion medium containing 3.2 g NaHCO_3_, 0.58 g L-glutamine, 0.11 g sodium pyruvate, 1.11 g HEPES, 1 bottle DMEM, 1 g RIA-grade BSA, and 70 mg ascorbate in 1 L ultra-purified water. The perifusion medium was filtered and de-gassed prior to preparation of secretagogues.

After retrieving islets from the perifusion chamber and placing them into a 60-mm non-treated culture dish, the IEQ of the retrieved islets was determined and islets were centrifuged for 3 min at 200 rcf at room temperature. Islet hormone extracts were prepared by incubating the retrieved islet pellet in 200 μL of fresh acid ethanol (50 μL 5N HCl + 5.5 mL 95% ethanol) for 24 hours. Samples were then spun at 3000 rcf for 5 min, aliquoted to 2 mL screwcap tubes, and stored at –80 °C until further processing.

Insulin concentration in perifusates and islet extracts was measured via ELISA (2020 – present; Human Insulin ELISA Kit, Mercodia, catalog # 10-1113-10) or RIA (Human Insulin-Specific RIA, Millipore catalog # HI-14K). In the same samples, glucagon was also measured via ELISA or RIA (2021 – present: Quantikine Glucagon ELISA Kit, R&D Systems catalog # DGCG0; 2020 – 2021: HTRF Glucagon Detection Kit, Cisbio catalog #62CGLPEH; prior to 2020: Glucagon RIA, Millipore catalog #GL-32K). For each assay, standards and quality controls were measured in duplicates, and perifusate and islet extract samples were measured in duplicates (RIA) or single point (ELISA). Secreted insulin and glucagon concentrations were normalized to IEQs and expressed as ng/100 IEQs/min. Alternatively, secreted insulin and glucagon concentrations were normalized to their respective total hormone content and expressed as %content/min.

#### Islet composition

To assess islet composition and acinar cell components, an aliquot of the human islet preparation was immobilized in a collagen I gel, embedded, and sectioned for histologic analysis. An aliquot containing approximately 500 IEQs of the human islet preparation was transferred into a 1.5 mL tube and washed three times in 1X PBS via centrifugation at 200 *xg* for 3 min. After removing the supernatant, 150 μL of collagen I working solution (563 μL collagen I stock, 167 μL ultrapurified H_2_O, 200 μL 5X DMEM, 20 μL HEPES, and 50 μL NaHCO_3_) was added to the islet pellet and the mixture was transferred into a well in the center of a 96-well plate. The plate was incubated at 37 °C for 1.5 hours to allow the gel to solidify, then fixed on ice for 15 min, filling the well with cold 4% paraformaldehyde/PBS. After the brief fixation, the paraformaldehyde solution was removed and, using a 28 G needle, the gel was loosened from the sides of the well and transferred with a spatula into a 12-well plate containing cold 4% paraformaldehyde/PBS. The gel was fixed for an additional 45 min, shaking on ice, before being washed three times with 1X PBS for 20 min shaking on ice, transferring the gel to a new well containing fresh 1X PBS for each wash. Following an overnight incubation in 30% sucrose/1X PBS, the collagen I gel was embedded into cryomolds containing OCT, imaged, and flash frozen on dry ice. Islet gels were stored at –80 °C until sectioning.

Following sectioning of collagen gels, immunofluorescence staining of islets was performed. In brief, 8-μm islet cryosections were allowed to thaw at room temperature and air-dried for 30 mins. Sections were washed with 1X PBS 3 times for 5 mins to remove the OCT, permeabilized with 0.2% Triton for 15 mins at room temperature, and washed 3 times in 1X PBS. Sections were blocked with 5% normal donkey serum in 1X PBS at room temperature for 90 mins in a humidified chamber, then incubated overnight at 4 °C in a humidified chamber with primary antibodies diluted in 0.1% Triton-X-100/1% BSA/1X PBS. Primary antibodies included: C-peptide (rat, Developmental Studies Hybridoma Bank, GN-ID4, RRID:AB_2631151, 1:100), Glucagon (mouse, Abcam, ab10988, RRID:AB_297642, 1:250), Glucagon (rabbit, Cell Signaling Technology, 2760S, RRID:AB_659831, 1:100), Somatostatin (goat, Santa Cruz Biotechnology, sc-7819, RRID:AB_2302603, 1:500), and HPX1 (mouse, Novus Biologicals, NBP1-18951, RRID:AB_1625456, 1:100). Sections were washed with 1X PBS 3 times for 10 mins each, then incubated for 1.5 hours at room temperature in a humidified chamber with secondary antibodies diluted in 0.1% Triton/1% BSA/1X PBS. Secondary antibodies were purchased from Jackson ImmunoResearch: Rat IgG-Cy2 (donkey, 712-225-150, RRID:AB_2340673, 1:500), Rat IgG-Cy5 (donkey, 712-175-150, RRID:AB_2340671, 1:200), Mouse IgG-Cy3 (donkey, 715-165-150, RRID:AB_2340813, 1:500), Rabbit IgG-Cy5 (donkey, 711-175-152, RRID:AB_2340607, 1:200), Goat IgG-Cy5 (donkey, 705-175-147, RRID:AB_2340415, 1:200), and Mouse IgG-Cy3 (donkey, 715-165-150, RRID:AB_2340813, 1:500). Sections were counterstained with 1:25,000 DAPI/PBS for 10 mins at room temperature, then washed 3 times for 15 mins each in 1X PBS. Sections were mounted with SlowFade Gold mounting medium, and islet sections were imaged using a high-resolution whole slide scanning system (ScanScope FL, Aperio/Leica) connected to a web-based digital slide repository powered by eSlide Manager and housed in the Vanderbilt University Medical Center Data Center. A quantitative assessment of islet cell composition and endocrine/acinar cell compartments was performed using a tissue classifier algorithm (Halo™, Indica Labs) to analyze 50-100 islets per labeling experiment. Images presented herein are deposited in the publicly available Pancreatlas™ platform^52^, which allows the exploration of full-resolution islet imaging data in an interactive manner (https://pancreatlas.org/datasets/853/explore).

#### Genetic analysis

DNA on 246/299 donors included in this release was isolated from human islet donor acinar tissue at Vanderbilt using the Wizard Genomic DNA Purification Kit (Promega, catalog # A1120). Quantification and quality assessment was performed using the Genomic DNA ScreenTape assay. DNA was normalized to 50 ng/μL prior to shipment to the HIGI for further analysis. For 30 samples no acinar tissue was available. For 9 of these, DNA was extracted from FPPE sections at Stanford using the Zymo QuickDNA FFPE Extraction Kit. For the remaining 23 donors DNA was isolated from acinar tissue at Stanford using the DNAExtraction (Qiagen DNeasyBlood&TissueKit, catalog # 69504).

Prior to genotyping all DNA samples were re-quantified using both the Nanodrop and Qubit platforms. Samples containing less than 500 ng total DNA were speed-vacuumed and concentrated prior to genotyping. All samples were genotyped using the Illumina Infinium Omni2.5Exome V1.5 array with ∼2.6 million single nucleotide variant (SNV) sites measured. Genotyping data underwent quality control by filtering of SNVs with excess missing genotypes (>2%), filtering of SNVs that do not conform to Hardy-Weinberg equilibrium (p<1×10^-6), and removal of samples with either excess missing genotypes (>3%) or discordant genetic vs recorded sex. For donor samples with excess missing genotypes, 10 μm-thick formalin-fixed paraffin-embedded pancreas tissue slides were requested from IIDP. DNA was extracted using the Zymo QuickDNA FFPE Extraction Kit and quantified by Nanodrop and Qubit. Samples were then restored using the Infinium HD FFPE Restore Protocol and then re-genotyped. Since FFPE samples have considerable DNA degradation, if these samples did not pass restoration, thicker 20 um FFPE slides were shipped from IIDP and DNA was extracted and sent for restoration and genotyping. For samples that had discordant sex, an XY PCR was performed on the original sample and an additional donor sample from IIDP (10-μm thick formalin-fixed paraffin-embedded pancreas tissue slides). Genotyping results from the 270 donors passing QC coordinates were then aligned to the positive strand on genome build GRCh38 and the data imputed against the TOPMed R3 reference panel. After quality control and imputation, 69.4 million high-quality SNVs (R2 > 0.3) and 269 samples were available for analysis. Genetic risk scores (GRS) for T1D and T2D were generated from previously published models using 67 and 338 SNVs respectively^7,41^. T2D partitioned genetic risk scores (pGRS) were generated using 94 SNVs with predetermined weights from a previous soft-clustering^42^. Genetic ancestry was determined using a random forest classifier trained on the first 20 principal components of the 1000 Genomes reference panel (https://github.com/gloynlab/GeneticAncestry).

#### Differentiational expression of predicted T2D effector transcripts in δ cells

Re-processed scRNA-seq data for islets of organ donors without diabetes were downloaded from PanKbase (https://pankbase-data-v1.s3.us-west-2.amazonaws.com/analysis_resources/single_cell_objects/min.cells0.01pct_min.features5pct_r mDoublets_harmony_data.Rds). The dataset included 87,505 islet cells (1877 δ cells) from 44 donors, https://zenodo.org/records/15588240, and is available for the lookup on Pancreatlas https://pancreatlas.org/datasets/1138/explore/omics/17. Additional lookup data for δ gene expression in human islets from donors without diabetes is available here https://powersbrissovalab.shinyapps.io/scRNAseq-Islets/. The variants (N = 338) in the calculation of the genetic risk score were mapped to the closest genes (N = 303) using BEDTools with the gene annotation from Ensembl (v112). The number of expressed genes in islets and δ cells was estimated by applying multiple thresholds, 0.001, 0.01, and 0.1, on the mean normalized counts across all genes and δ cells only, respectively. Differential gene expression analysis between the δ cells and the rest of the cells, and each of the other cell types, on the closest genes was performed using the Wilcoxon rank-sum method in the scanpy package (v1.10.4), with p values adjusted by the Benjamini-Hochberg method. Significant genes (padj < 0.05) that are upregulated in δ cells (logfoldchanges > 0, higher expression in δ cells than in other types of cells) and expressed in at least 20% of δ cells are visualized in dot plot showing mean gene expression and fraction of cells, ordered by descending logfoldchanges, and heatmap showing effect size (logfoldchanges), with the marker gene of δ cells (SST) included for reference. The blue color in **Figure 7F-G** highlights genes with a fold change greater than 2 when comparing δ cells with the remaining cells. A Fisher test was performed to check if there is an enrichment of the closest genes in the differentially expressed genes between δ cells and other cell types.

### QUANTIFICATION AND STATISTICAL ANALYSIS

#### Hormone secretion traits

Using the insulin and glucagon secretion traces normalized to IEQ, eleven insulin secretion traits and nine glucagon secretion traits were defined and calculated as illustrated in **Figure 2C-D**.

#### Statistical analysis

To determine demographic, islet processing, and morphologic variables associated with differences in islet secretory function, we first performed Spearman correlation analyses. For variables with a statistically significant association with islet secretory traits, we utilized multivariable regression models to examine the relationship among the explanatory variables and the outcome variable of interest. For each model, we controlled for the following potential confounders, unless otherwise noted: donor age, sex, BMI, isolation center, HbA1c, reported race or ethnicity, pre-shipment culture time, and transit time. A statistically significant regression coefficient for the explanatory variable (p-value < 0.05) denoted a significant association. Given multiple p-values, we adjusted for the false discovery rate (FDR) at 0.05 to control for multiple comparisons.

Similarly, to explore potential differences based on prediabetes status or sex, we used multivariable regression models. In these models, an indicator variable (0 for normal, 1 for pre-diabetes participants; 0 for male, 1 for female) acted as the explanatory variable. For prediabetes-related associations, we controlled for all eight of the aforementioned potential confounders except HbA1c. For sex-related associations, we controlled all potential confounders except donor sex. The significance of the regression coefficient (b) for the group indicator (p-value < 0.05) indicated a notable difference in the outcome variable between the two groups.

To ensure comparability of regression coefficients across models, we normalized the outcome variables and all continuous explanatory variables, including covariates. This normalization involved centering the variables by subtracting their mean and scaling them by their standard deviation. However, this procedure was not applied to categorical covariates in the model, such as center, self-reported ancestry, and sex. Statistical analyses were all performed using R Statistical Software version 4.3.0 (R Foundation for Statistical Computing, Vienna, Austria) and GraphPad Prism version 10.

