## Supplemental File 1 for "Heterogeneous endocrine cell composition defines human islet functional phenotypes"

**IIDP Coordinating Center, City of Hope:** Martha Antler, Jenny Chuang, James Cravens, Jessica Girard, Julie Hom, Julie Kilburn, Barbara Olack, Marcelo Pineda, Dajun Qian, Heather Sibley, Janice Sowinski, Shaunna Spears, Carol Swanson, Julia Yousefi

**Columbia University Islet Core**: Bhawna Chandravanshi, Xiaojuan Chen, Alexey Koshkin, Stuart Weisberg

**Human Islet Core Facility at Loyola Medical Center**: Mehboob Ali, Casi Blanton, Luis Fernandez, Karol Palasiewicz, Yucel Yankol, Aparna Yellapa

**Imagine Islet Center, Imagine Pharma**: Rita Bottino, James Eskra, Yesica Garciafigueroa

**Nationwide Children's Islet Transplant & Biology Facility:** Balamurugan N. Appakalai, Ahad A. Kodipad, Chandrashekar B. Revanna, Krishna K. Samaga

**Prodo Labs Human Islet Isolation Center**: Gowri Arulmoli, David Boundurant, Dylan Cardiff, Nora Dawood, Tiffany Enriquez, Brian Haight, Ergeng Hao, Virginia Keach, Daniel McRaney, Sally Padayao, Rebekah Poblete, Mark Ramon, Stefan Riedel David Scharp, Matt Wright

**Southern California Islet Cell Resource Center, City of Hope**: Tania Aguilar, Ismail Al-Abdullah, Alina A Avakian, Shiela Bilbao, Randi Collins, Mohamed Elsayed, Kevin Ferreri, Pablo Garcia, Debbie Gish, Noe Gonzales, Lam Ho, Itzia Iglesias-Meza, Fouad Kandeel, Alexander Kaye, Willem M Kühtreiber, Doreen Ligot, Leonard Medrano, Miryam Mehra, Aria Miller, Alina Oancea, Keiko Omori, Meirigeng Qi, Karen Ramos, Jeffrey Rawson, Mayra Salgado, Amanda Serrano, Jeannette Stratton, Autumn Tate, Jeffrey Tauer, Ivan Todorov, Amber Tucker, Luis Valiente, Robin Zafra, Emily Zebadua

**University of Arizona, Institute for Cellular Transplantation**: Jose Cano, Carola Davila, Trisha Marie Fabijanic, Stacy Hyde, Chan Ion, Amy Kelly, Klearchos K Papas, Craig Weber

**University of Miami**: Rodolfo Alejandro, Alex Alvarez, Carmen Castillo, Maxwell Donaldson, Ross Haertter, Omaima Hanif, Itzia Iglesias, Aisha Khan, Clarissa Lenero, Elina Linetsky, Eric M LoRusso, Jorge Montelongo, Kevin Peterson, Camillo Ricordi, Rudy U Rodriguez, John Sotolongo, Joel Szust, Xiao Jing Wang, Xiumin Xu,

**University of Pennsylvania Islet Transplant Center**: Insuk Choe, Yanjing Li, Chengyang Liu, Diane McLaughlin, Zaw Min, Ali Naji, Roman Prosniak, Johanna Sotiris, Jing Wang, Min Wang, Wei Wang, Amy Wexler, Xi Zuo

**University of Wisconsin Human Islet Core**: Casi Blanton, Connie Chamberlain, Peter Chlebeck, Sebastian Danobeitia, Michael Eerhart, Luis Fernandez, Ayesha Khan, Xiaobo Ma, Jon Odorico, Jose A Reyes, Sara Sackett, Ellen Abad Santos, Tiffany Zens, Martynas Ziemelis, Laura Zitur

**VCU Islet Cell Processing Lab**: Elijah Burch, Jagan Kalivarathan, Mazhar A Kanak, Shujauddin Mohammed, Prathab B Saravanan, Yoshiko Tamura
