## Supplementary material for "Heterogeneous endocrine cell composition defines human islet functional phenotypes": Extanded Data Figures and Tables

### Extended Data Figure 1

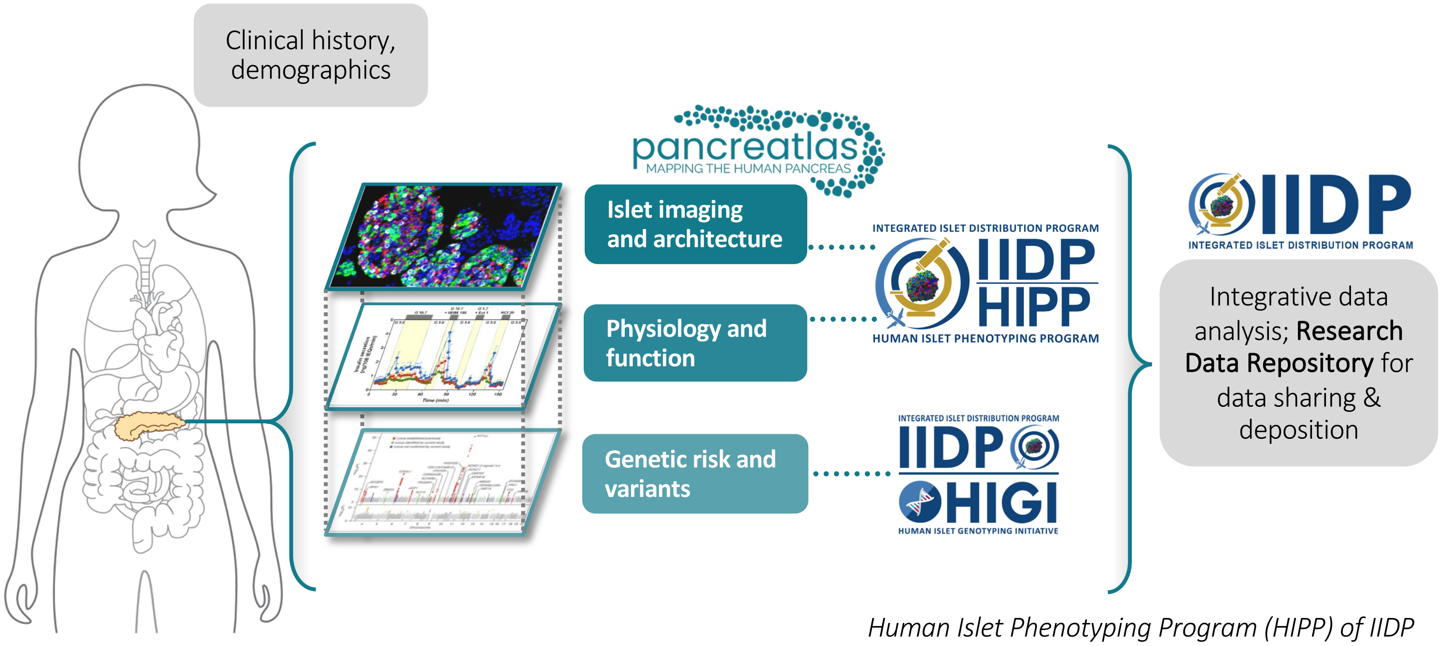


*Human Islet Phenotyping Program (HIPP) of IIDP*

*Human Islet Genotyping Program (HIGI) of IIDP*

Extended Data Figure 2

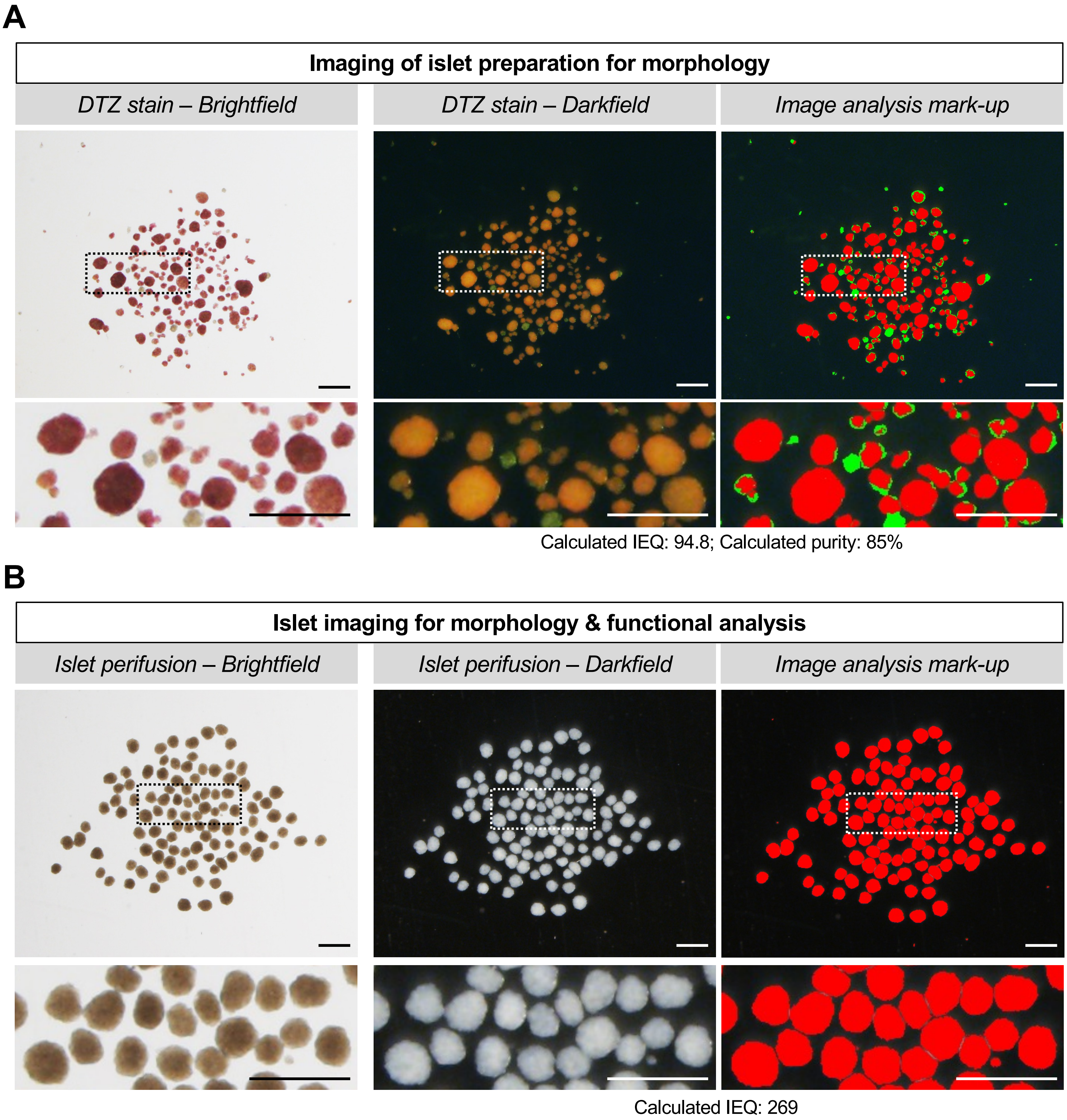

### Extended Data Figure 3

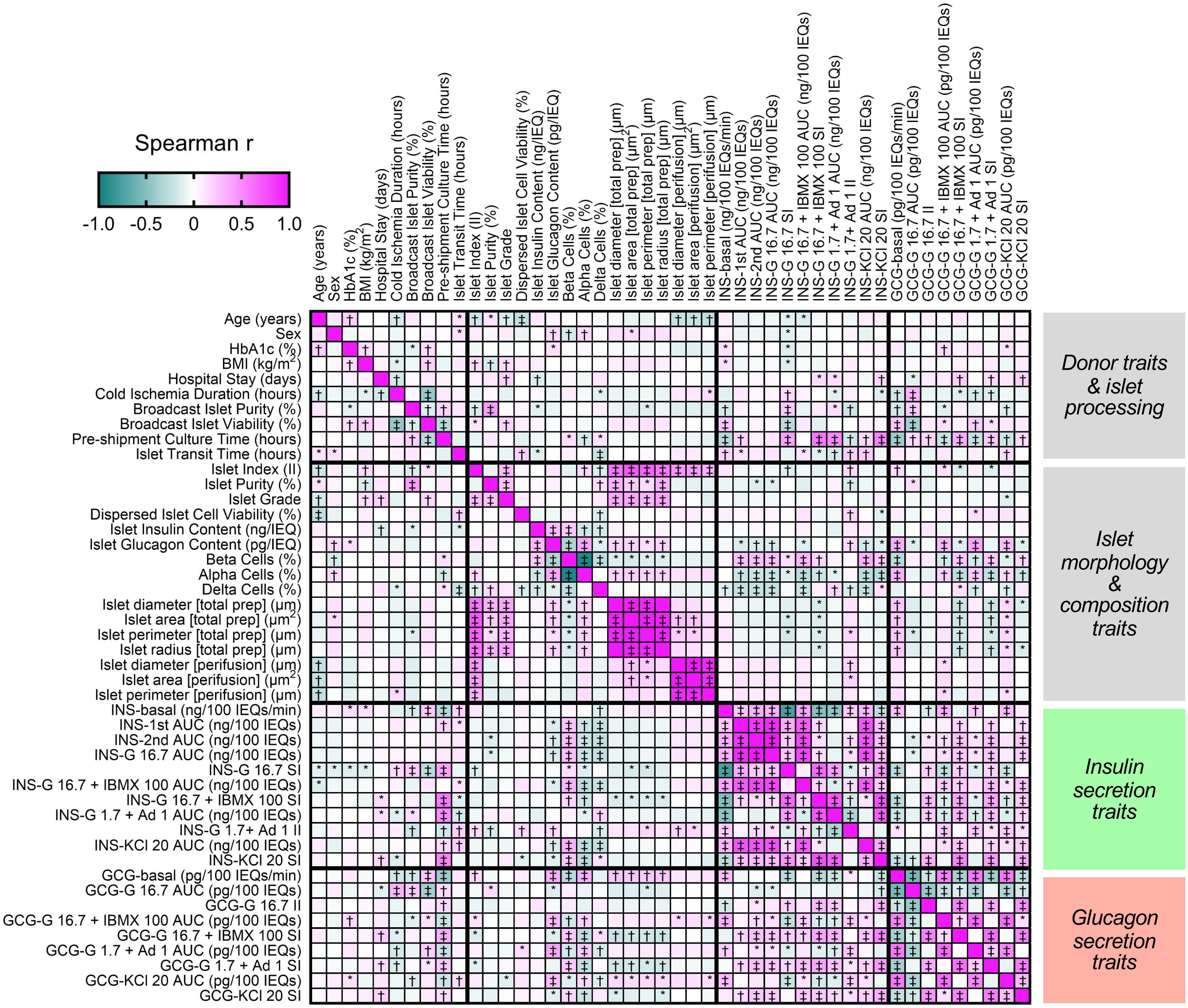

### Extended Data Figure 4

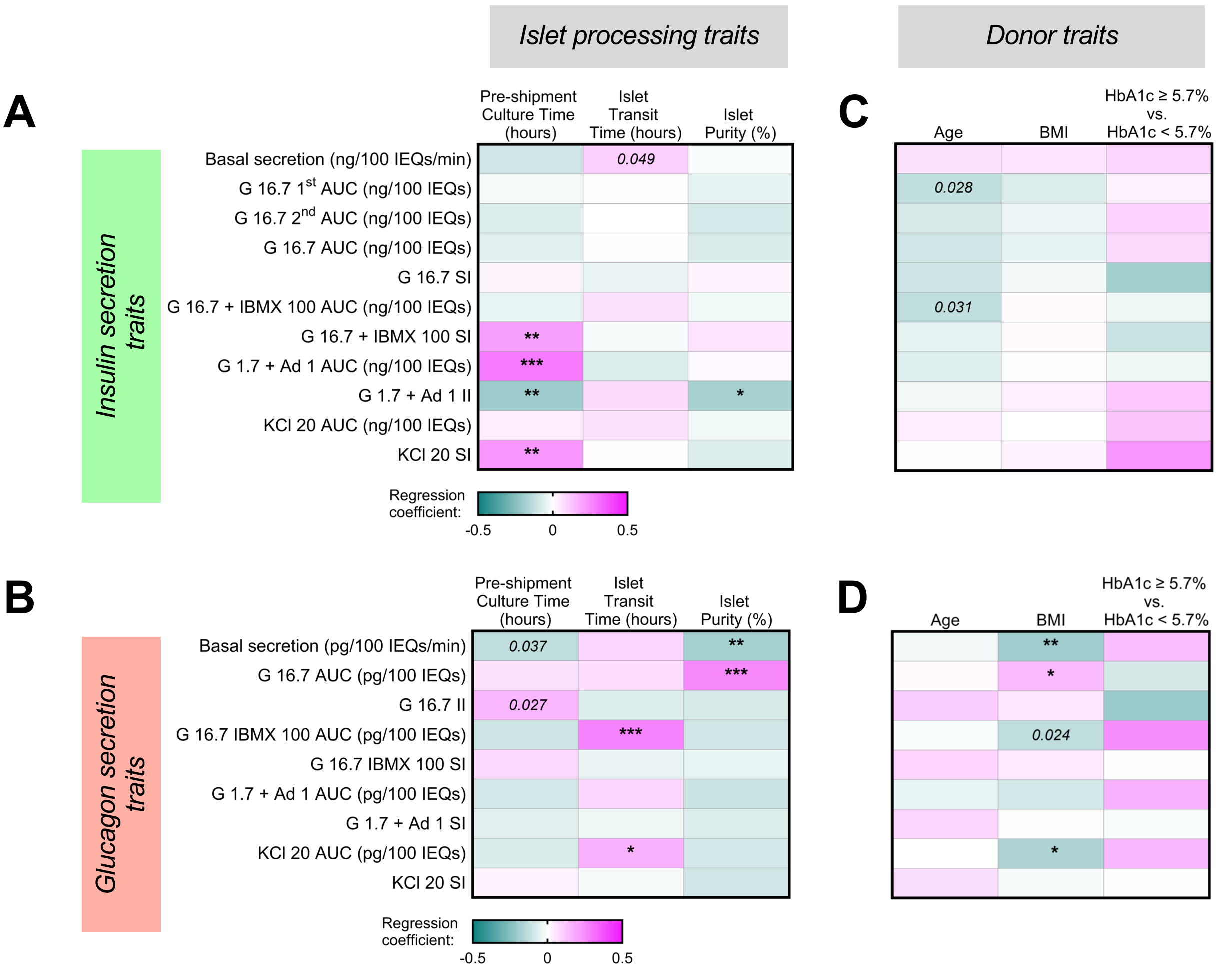


**E**

| Insulin secretion traits | | Basal secretion (ng/IEQ) | G 16.7 1st AUC (ng/IEQ) | G 16.7 2nd AUC (ng/IEQ) | G 16.7 AUC (ng/IEQ) | G 16.7 SI | G 16.7 + IBMX 100 AUC (ng/IEQ) | G 16.7 + IBMX 100 SI | G 1.7 + Epi 1 AUC (ng/IEQ) | G 1.7 + Epi 1 II | KCl 20 AUC (ng/IEQ) | KCl 20 SI |
|---|---|---|---|---|---|---|---|---|---|---|---|---|
| Global adjusted p-value *(unadjusted p-value)* | Sex | n.s. | n.s. | n.s. | n.s. | n.s. | n.s. | n.s. | n.s. | n.s. | n.s. | n.s. |
| | Reported race or ethnicity | 0.0671 *(0.012)* | n.s. | n.s. | n.s. | n.s. | 0.103 *(0.037)* | 0.103 *(0.036)* | n.s. | n.s. | 0.0671 *(0.011)* | n.s. |
| | Genetic Ancestry | 0.045 | n.s. | n.s. | n.s. | n.s. | n.s. | n.s. | n.s. | n.s. | 0.1482 *(0.027)* | n.s. |
| | Isolation Center | 2.46E-06 | 1.30E-05 | 3.66E-04 | 1.15E-04 | 4.78E-08 | 1.68E-03 | 1.49E-02 | 7.95E-05 | 2.08E-02 | 1.30E-03 | 7.20E-03 |

**F**

| Glucagon secretion traits | | Basal secretion (pg/IEQ) | G 16.7 AUC (pg/IEQ) | G 16.7 II | G 16.7 IBMX 100 AUC (pg/IEQ) | G 16.7 IBMX 100 SI | G 1.7 + Epi 1 AUC (pg/IEQ) | G 1.7 + Epi 1 SI | KCl 20 AUC (pg/IEQ) | KCl 20 SI |
|---|---|---|---|---|---|---|---|---|---|---|
| Global adjusted p-value *(unadjusted p-value)* | Sex | n.s. | n.s. | n.s. | n.s. | n.s. | n.s. | n.s. | n.s. | 0.31 *(0.034)* |
| | Reported race or ethnicity | n.s. | n.s. | n.s. | n.s. | n.s. | n.s. | n.s. | n.s. | n.s. |
| | Genetic Ancestry | n.s. | n.s. | n.s. | n.s. | n.s. | n.s. | n.s. | n.s. | n.s. |
| | Isolation Center | 1.01E-06 | 6.37E-09 | 3.28E-03 | 1.05E-02 | n.s. | 8.66E-04 | n.s. | 3.28E-03 | n.s. |

### Extended Data Figure 5

*n* = 86 Admixed American, *n* = 21 African, *n* = 11 East Asian, *n* =150 European

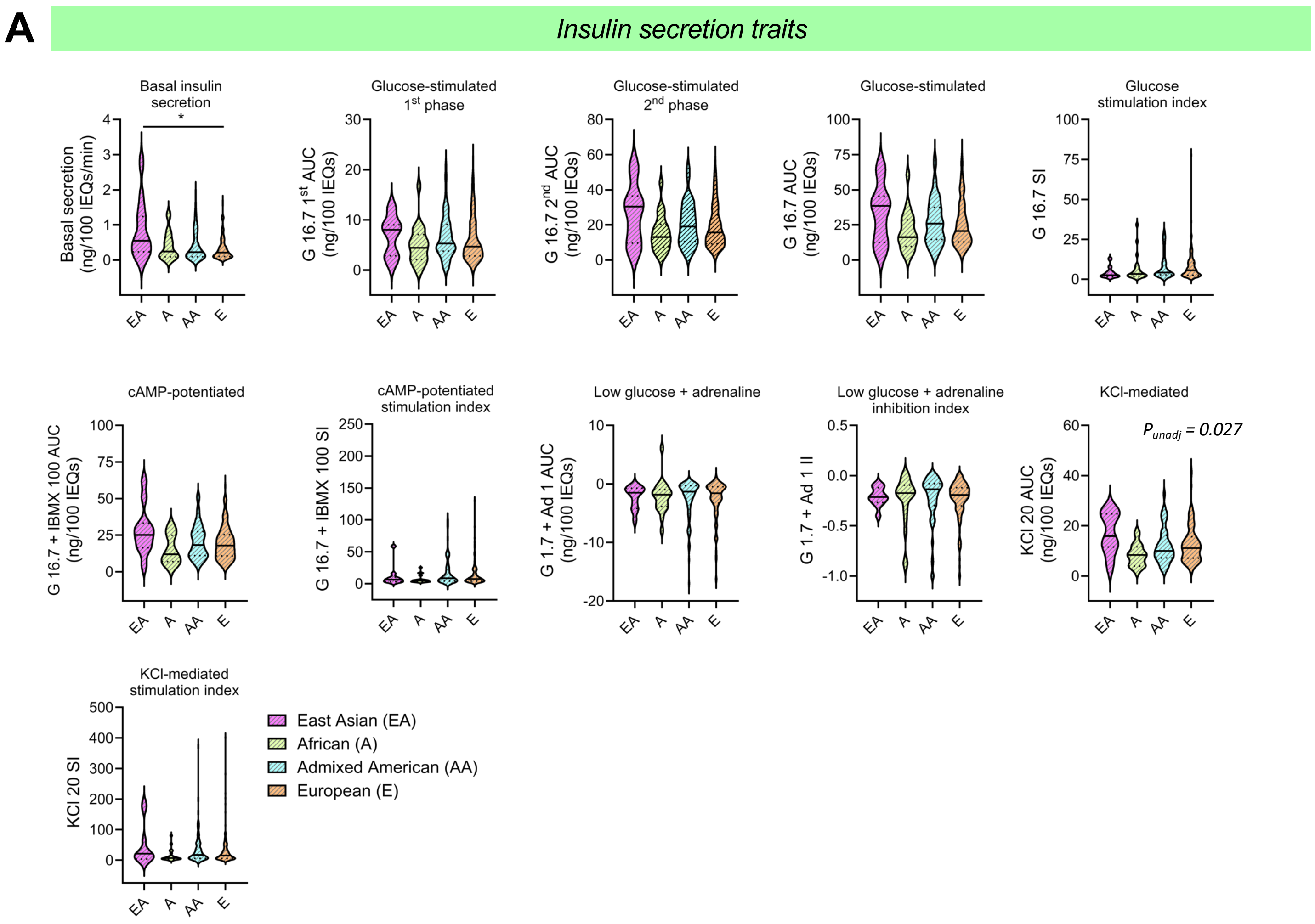


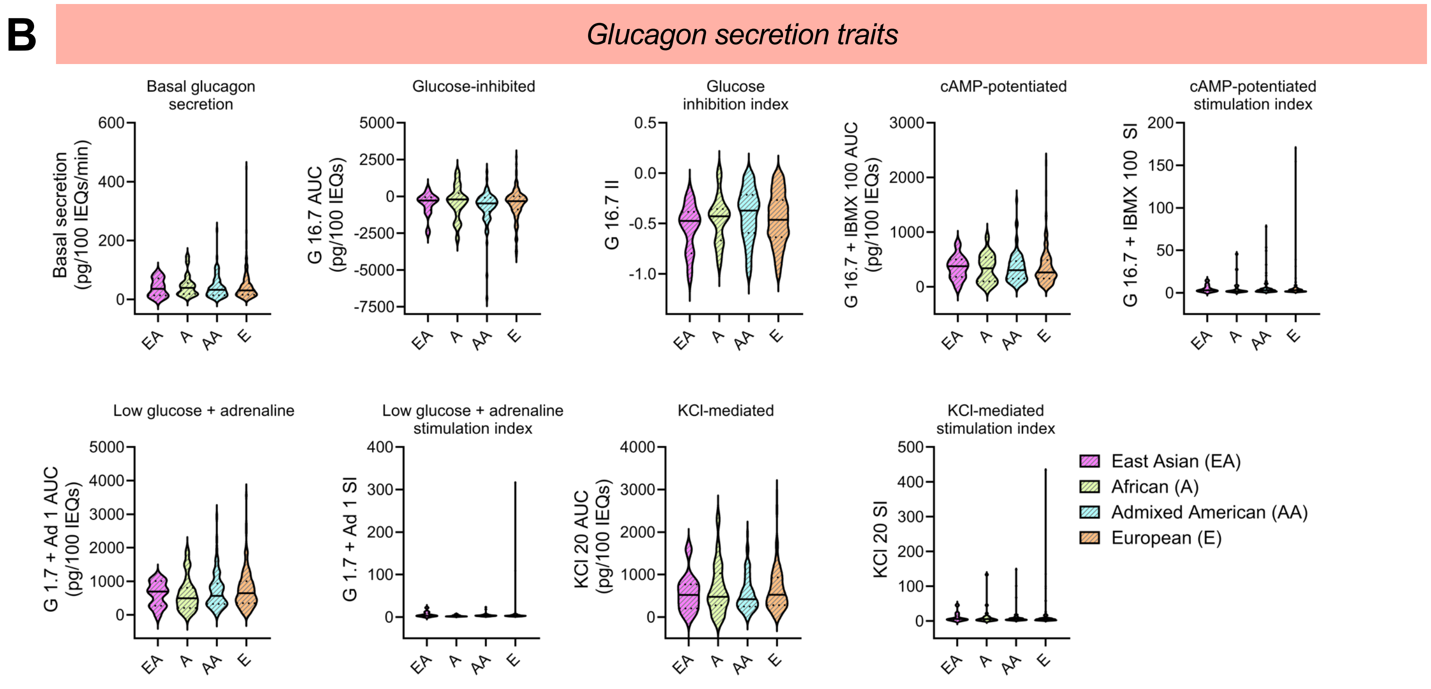

### Extended Data Figure 6

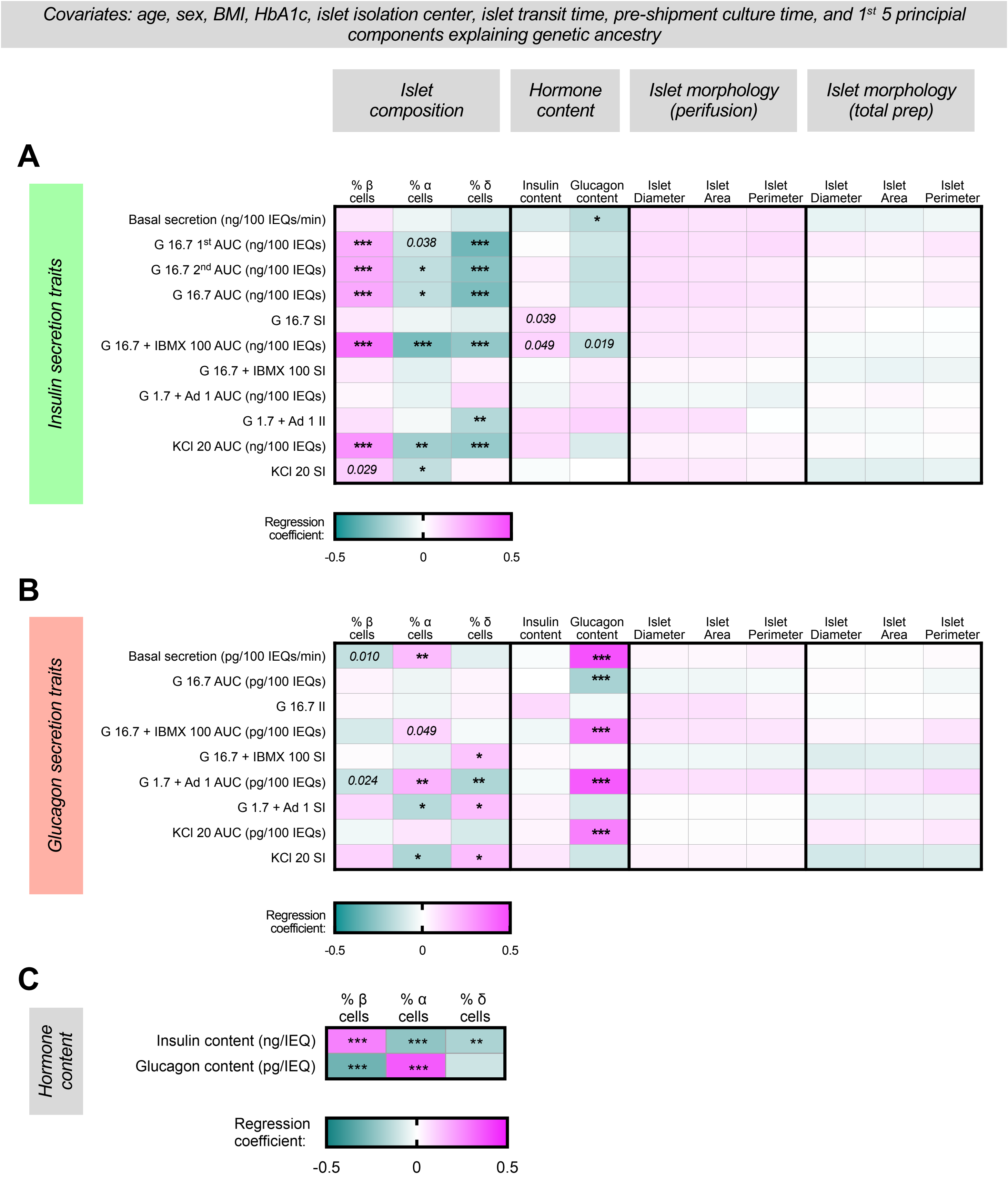

### Extended Data Figure 7

**A**

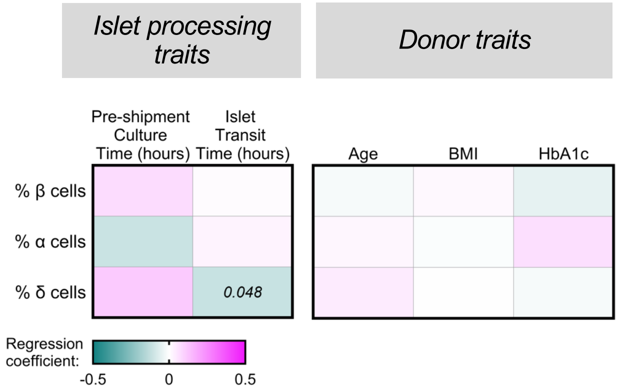


**B**

| Islet composition | | % β cells | % α cells | % δ cells |
|---|---|---|---|---|
| Global adjusted p-value | Sex | 0.0065 | 0.007 | n.s |
| | Reported race or ethnicity | 0.003 | 0.003 | n.s |
| | Genetic Ancestry | 0.0004 | 0.0004 | n.s |
| | Isolation Center | n.s. | n.s. | n.s. |

**C**

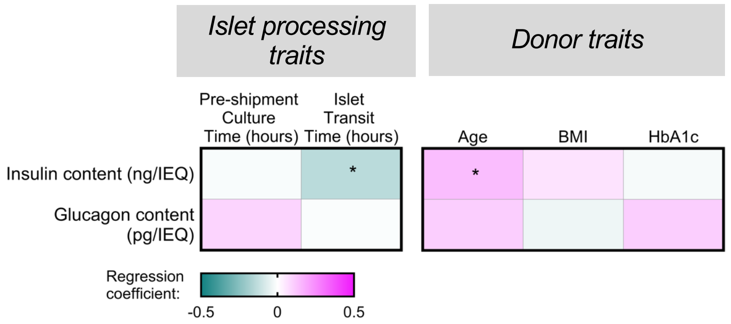


**D**

| Hormone content | | Insulin content (ng/IEQ) | Glucagon content (pg/IEQ) |
|---|---|---|---|
| Global adjusted p-value | Sex | n.s. | 0.015 |
| | Reported race or ethnicity | n.s. | n.s. |
| | Genetic Ancestry | 0.045 | n.s. |
| | Isolation Center | n.s. | n.s. |

*Covariates: age, sex, BMI, HbA1c, islet isolation center, islet transit time, pre-shipment culture time 1st 5 principial components explaining genetic ancestry*

**E**

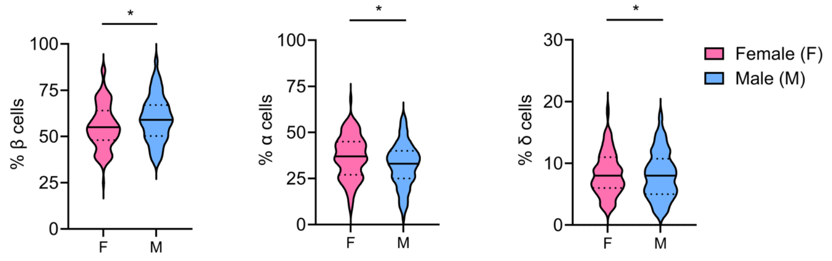


**F**

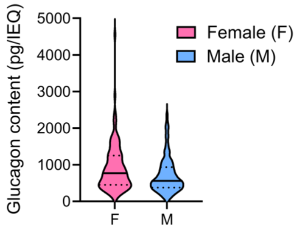

### Extended Data Figure 8

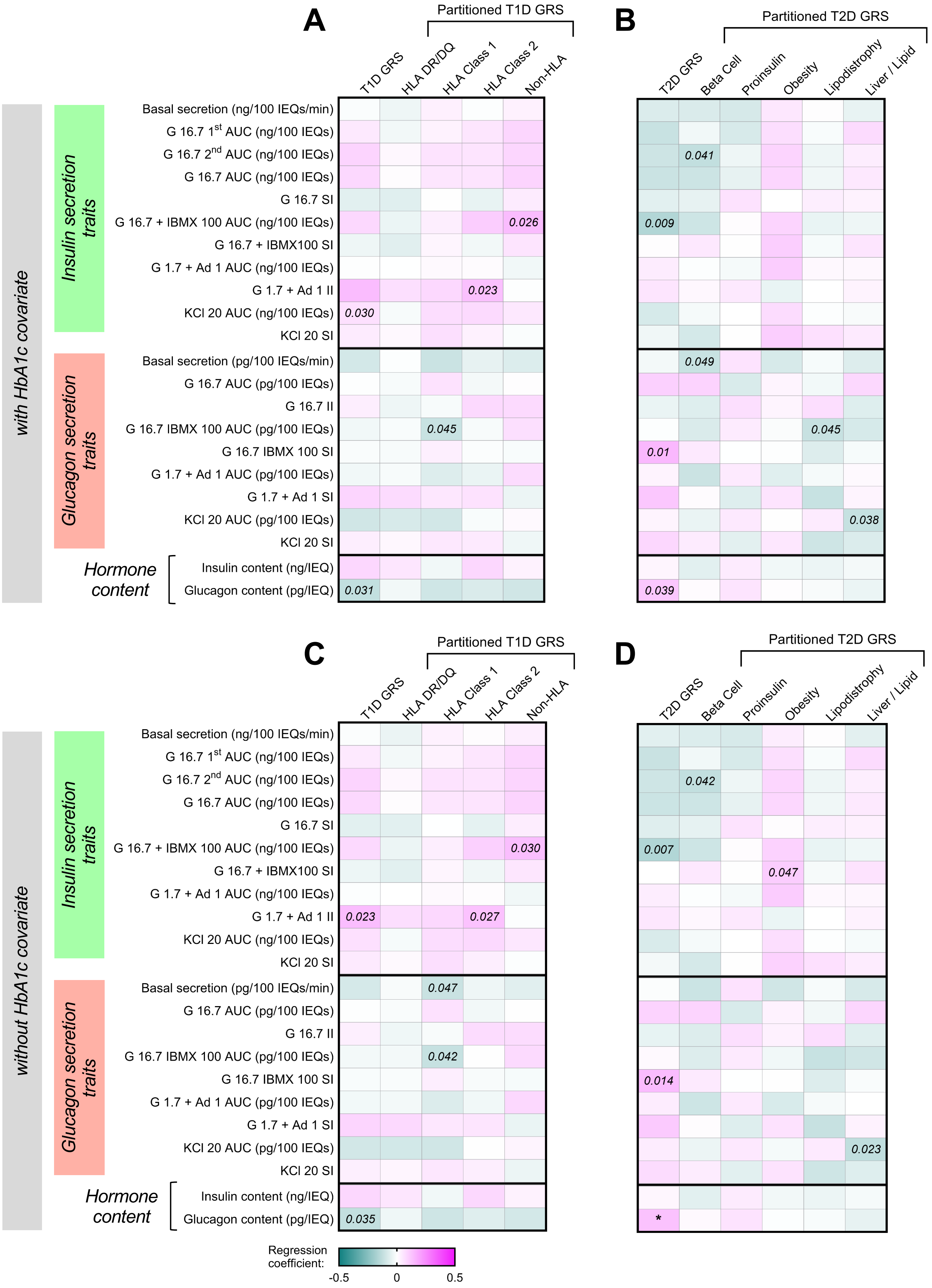

**Extended Data Figure 9**

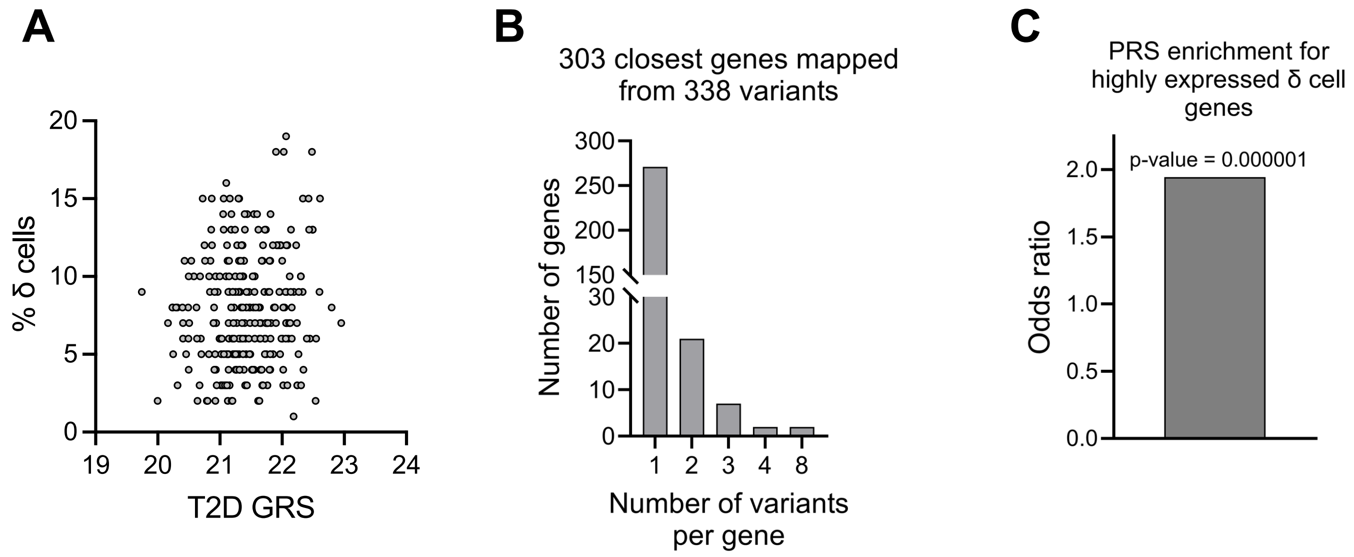
